# The 5-HT1F Receptor Agonist Lasmiditan improves Cognition and Ameliorates Associated Cortico-Hippocampal Pathology in Aging Parkinsonian Mice

**DOI:** 10.1101/2025.02.13.638147

**Authors:** Atsushi Ishii, Jasmine R Meredith, Mandi J Corenblum, Kelsey Bernard, Paige V Wene, Nainika Menakuru, Paul V Santiago, Rick G Schnellmann, Lalitha Madhavan

## Abstract

While the etiopathology of Parkinson’s disease (PD) is complex, mitochondrial dysfunction is established to have a central role. Thus, mitochondria have emerged as targets of therapeutic interventions aiming to slow or modify PD progression. We have previously identified serotonergic 5-HT1F receptors as novel mediators of mitochondrial biogenesis (MB) - the process of producing new mitochondria. Given this, here, we assessed the therapeutic potential of the FDA-approved 5-HT1F receptor agonist, lasmiditan, in a chronic progressive PD model (Thy1-aSyn ‘line 61’ mice). It was observed that systemic lasmiditan exhibited robust brain penetration and reversed cognitive deficits in young (4-5.5 months old) Thy1-aSyn mice (1mg/kg, every other day). Anxiety-like behavior was also improved while motor function remained unaffected. These behavioral changes were associated with enhanced MB and mitochondrial function, paired with reduced alpha-synuclein aggregation particularly in cortico-hippocampal regions. Furthermore, in older (10-11.5 months old) mice, although the effects were milder, daily lasmiditan administration increased MB and bettered cognitive abilities. In essence, these findings indicate that repurposing lasmiditan could be a potent strategy to address PD-related cognitive decline.

## INTRODUCTION

Parkinson’s disease (PD) is the fastest growing neurological disorder worldwide, with a prevalence set to double by 2040^1,2^. In addition to the characteristic motor symptoms, multiple non-motor symptoms occur in PD that contribute greatly to the overall disease burden. Cognitive impairment is a key non-motor symptom which is known to be integral to the natural history of the disease. It can potentially occur at any disease stage and is known to be up to six times more common in individuals with PD than in the healthy population^3–6^. While symptomatic treatments that can provide transient relief, especially in the context of movement problems, are available, there are no therapies that address cognitive issues or broadly alter the course of the disease.

Overall, PD is understood to have a complex multi-factorial etiology involving interactions between several genetic and environmental factors over the course of aging^5^. Although these gene-environment interactions are not fully understood, studies have demonstrated a crucial and early role of mitochondrial dysfunction in PD^7–11^. In fact, the failure of mitochondrial quality control is a well-known widespread feature in both sporadic and genetic forms of PD^7–12^, and can contribute to several pathological cascades implicated in PD^13–16^. An important pathological mechanism involves the bidirectional interaction between mitochondrial stress and alpha-synuclein (α-synuclein) aggregation. For instance, the accumulation of α-synuclein in mitochondria can reduce complex I activity, increase reactive oxygen species, inhibit mitochondrial protein transport and reduce mitochondrial function^17,18^. On the other hand, mitochondrial dysfunction can promote oxidative stress, damaging cellular and vesicular membranes, and lead to the α-synuclein aggregation itself^17,18^.

Given the central role that mitochondria play in PD, they have naturally emerged as a therapeutic target. Agents which have shown promise in preclinical studies include creatine, vitamin E, mitoquinone, genistein, and resveratrol among others^19–21^. These approaches mainly modulate mitochondrial dysfunction by controlling oxidative stress, improving complex I function or mitochondrial potential and inhibiting mitochondrial-induced apoptosis. However, these tactics address single aspects of mitochondrial dysfunction, and have largely been ineffective as none has proved to have an unequivocal disease-modifying effect in human clinical trials^13,19^.

Enhancement of mitochondrial biogenesis (MB) offers a more comprehensive way to improve mitochondrial function by improving multiple aspects of mitochondrial biology. MB is the process by which new mitochondria are formed within the cell and is crucial for maintaining cellular energy homeostasis and function^22^. Specifically, MB is a transcriptional program that involves an intricate network of pathways for both nuclear- and mitochondrial DNA-encoded genes. MB is governed by peroxisomal proliferator coactivator-1α (PGC-1α), the “master regulator of MB,” which controls the expression of this network^23^. PGC-1α interacts with and co-activates several transcription factors, including nuclear respiratory factors (Nrf) 1 and 2, resulting in the transcription of not only nuclear-encoded mitochondrial genes, but also mitochondrial transcription factor A (TFAM), which activates transcription of mitochondrial-encoded genes^23^. Nuclear-encoded proteins are then transferred to the mitochondria, where nuclear- and mitochondrial-encoded subunits of the electron transport chain are assembled. PGC-1α/TFAM are reduced following ischemic injuries in multiple organs^24–26^. In PD, impaired MB has been observed, leading to a decrease in mitochondrial mass and function^27^, which in turn exacerbates the vulnerability of neurons to oxidative stress and energy deficiency, ultimately contributing to neurodegeneration^5,28^.

Thus, in this study, we proposed to address PD-associated mitochondrial dysfunction using pharmacological enhancement of MB through 5-hydroxytryptamine receptor 1F (5-HT1F) agonism. We previously identified the 5-HT1F receptor as a novel mediator of MB^29,30^. In particular, we showed that treatment with the selective and potent 5-HT1F receptor agonist, LY344864, increased MB and 1) restored kidney function following ischemia/reperfusion injury, and 2) contusion-induced spinal cord injury^31–33^. We have also demonstrated in a 6-hydroxydopamine rat PD model that LY344864 rescues tyrosine hydroxylase neurons and improves locomotor activity via 5-HT1F receptor-mediated MB^8,34–37^ . In extension, the current study tests the therapeutic potential of lasmiditan, a US Food and Drug Administration (FDA)-approved 5-HT1F receptor agonist. Lasmiditan is approved by the FDA for the acute treatment of migraine headaches^38^, and is a high-affinity and highly selective 5-HT1F receptor agonist with a favorable safety profile, making it a promising therapeutic candidate^39,40^. Here, we utilize Thy1-aSyn (line 61) mice that overexpress full-length human wild-type α-synuclein, under the Thy1 promoter, to test lasmiditan efficacy.^41^ The Thy1-aSyn mice reproduce many features of early pre-motor, motor and non-motor aspects of sporadic PD in an age-relevant manner including associated pathological, biochemical, and molecular alterations. In a nutshell, using both young and aged Thy1-aSyn mice, we show that lasmiditan improves MB and mitochondrial function, reduces α-synuclein, and particularly improves cognitive symptoms.

## MATERIALS AND METHODS

### Animals

Hemizygous human Thy1-aSyn mice (ASyn) (C57BL/6-DBA/2 background) and wild-type (WT) mice were maintained in the University of Arizona Animal Care Facility^42–45^. Male mice were mainly used, given the milder phenotype in female mice due to the location of the transgene on the X chromosome that can get randomly inactivated^41,43^. Male littermates used for the study were housed in standard cages, and the genotypes of all mice were confirmed by PCR analysis of tail DNA. The mice were housed under a reversed 12-hour light-dark cycle, with food and water available ad libitum. Mice were handled according to the rules and regulations of the National Institutes of Health (NIH) and Institutional Guidelines on the Care and Use of Laboratory Animals, as well as the Animal Research: Reporting of In Vivo Experiments guidelines. All experimental protocols were approved by the University of Arizona Institutional Animal Care and Use Committee.

#### Experimental Design

ASyn mice at 4 or 10 months of age were administered 1 mg/kg of Lasmiditan (Achemblock, San Diego, CA, USA) via intraperitoneal (i.p.) injection every other day for 6 weeks (**Fig. 2A**). WT and ASyn mice of the same age were used as controls, receiving equivalent volumes of normal saline under the same administration schedule. An additional group was included with respect to the 10-month-old study, wherein mice were treated with 1 mg/kg of lasmiditan daily for 6 weeks via i.p. injection. Control mice for this group also received daily saline injections for 6 weeks. Behavioral experiments were initiated at the 4th week following the start of i.p. injections, and brain samples were collected at the end of the 6th week.

**Figure 1.**
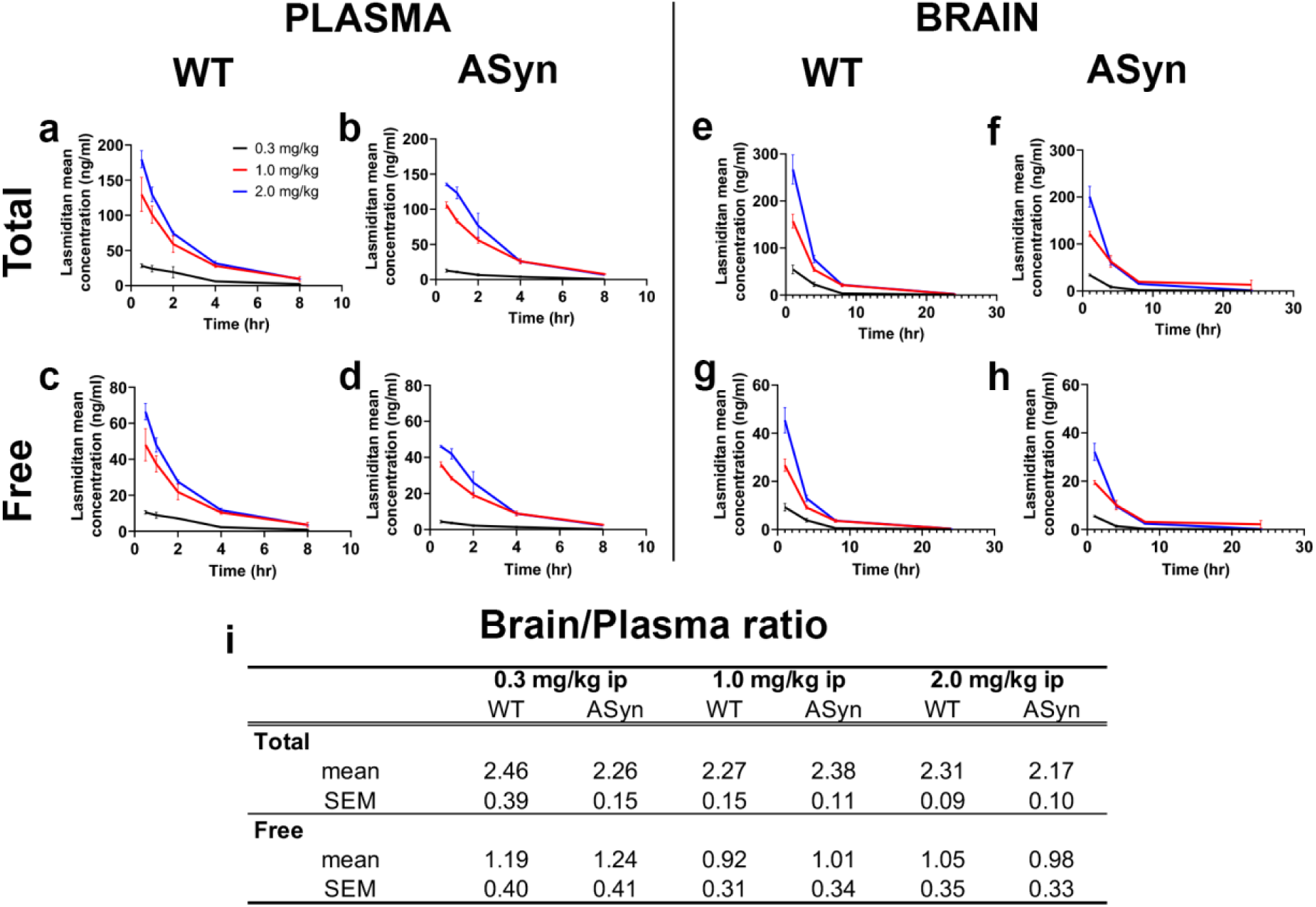
Pharmacokinetics of lasmiditan. This figure shows the time-dependent concentrations of total and free lasmiditan in the plasma and brain of WT and ASyn mice following a single intraperitoneal injection of 0.3 mg/kg (black line), 1.0 mg/kg (red line), or 2.0 mg/kg (blue line). Total lasmiditan concentrations in the plasma of WT and ASyn mice are shown in (a) and (b), respectively, while Free lasmiditan concentrations are shown in (c) and (d). Plasma levels were below the limits of detection at 24 hrs. Total lasmiditan concentrations in the brain of WT and ASyn mice are displayed in (e) and (f), and Free lasmiditan levels are in (g) and (h). (i) indicates the ratio of total and free lasmiditan concentrations between the brain and plasma at doses of 0.3, 1.0, and 2.0 mg/kg. N=3 mice/group, data are expressed as mean ± SEM.

**Figure 2.**
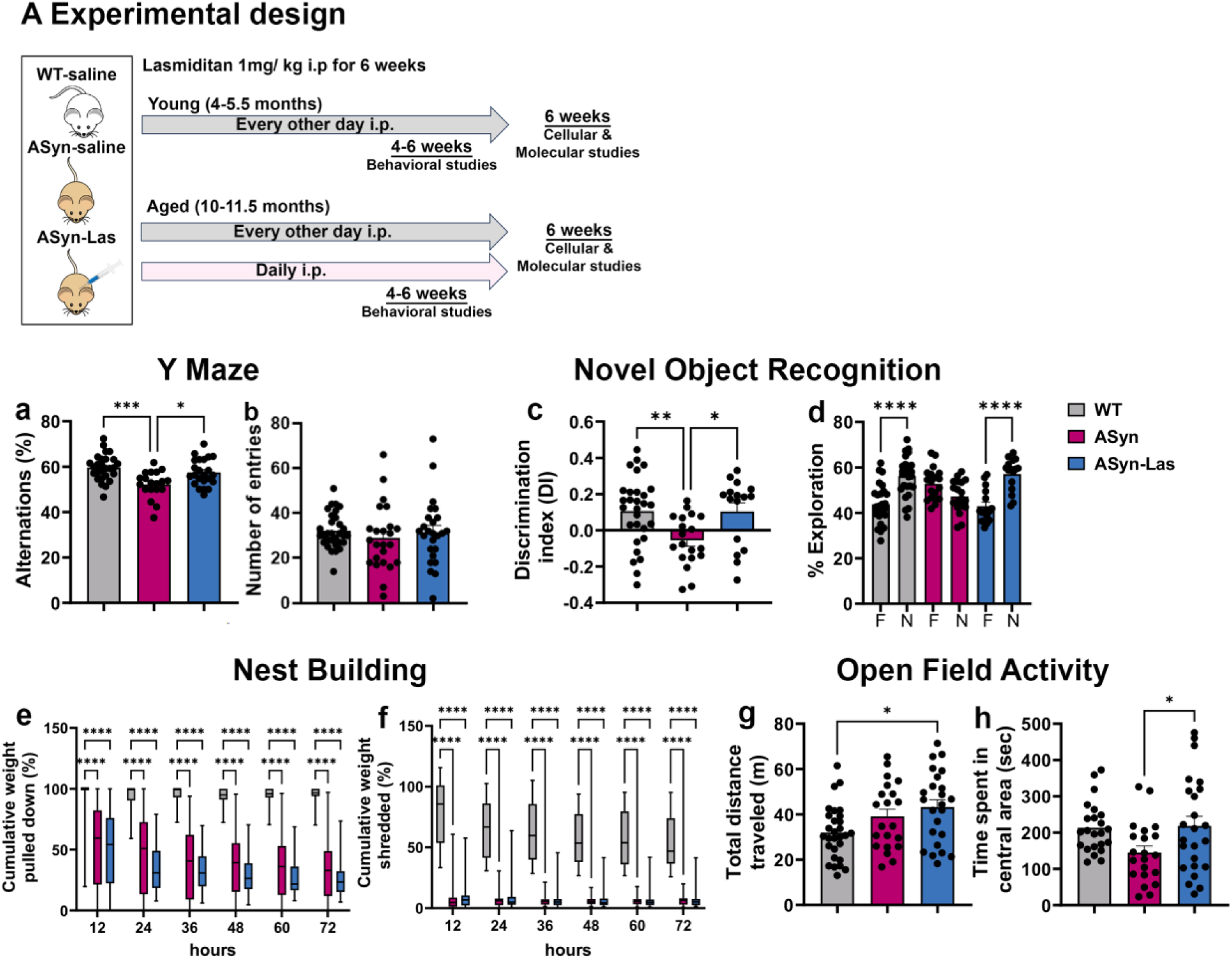
Experimental Design and behavioral assessment of young mice. (A) illustrates the overall experimental design of the studies involving young (4-5.5 mos old) and aged (10-11.5 months old) mice. WT: wild type; ASyn: Thy1-aSyn transgenic mice; ip: intraperitoneal injection; Las: lasmiditan. The effects of lasmiditan on motor and cognitive functions were evaluated. (a, b) displays results from a Y-maze task. WT (gray), ASyn (magenta), and ASyn-Las (blue) mice were compared with regards to alternation rates (a) and the number of arm entries (general activity levels, b). The mice were also subjected to a novel object recognition (NOR) test, results of which are expressed as a discrimination index (c, DI) and the percentage of time spent exploring the novel versus familiar object (d). A complex nest building test examined gross motor function and motivation by measuring the amount of cotton nestlets pulled down (e), and fine motor function by computing the amount of nestlets shredded (f). Locomotor activity and anxiety-like behavior were evaluated via an open field task. The total distance traveled, and the time spent in the center of the open field are displayed in (g) and (h) respectively. **** p < 0.0001, *** p = 0.001, ** p < 0.01, * p < 0.05; One-way ANOVA with Tukey’s tests for multiple comparisons, Two-way repeated-measures ANOVA, or mixed-effects model with Holm-Šidák’s tests for multiple comparisons; mean ± SEM, n = 16-31 animals/group.

### Lasmiditan

Lasmiditan was obtained from Advanced ChemBlock Inc (#22337, Hayward, CA, USA) and dissolved in Dimethyl Sulfoxide (DMSO: Sigma-Aldrich, St. Louis, MO, USA) to prepare a stock solution at a concentration of 10 mg/mL. The stock solution was further diluted with saline to a final concentration of 0.5 mg/mL. The diluted solution was administered intraperitoneally.

### PK study

WT (N=27) and ASyn (N=27) mice were divided into three groups to receive Lasmiditan at doses of 0.3, 1.0, or 2.0 mg/kg. Lasmiditan was administered via i.p., and blood plasma samples were collected at 0.5, 1, 2, 4, 8, and 24 hours post-injection (n=3 per time point per dose). In addition, brain tissues were collected at 4, 8, and 24 hours post-injection (n=3 per time point per dose) for further analysis. For both plasma (10 µL) and brain homogenate (20 µL) samples, 10 µL of internal standard (aza-lasmiditan) in acetonitrile was added, followed by protein precipitation with 30 µL acetonitrile. After vortexing, samples were frozen for 10 minutes, then centrifuged at 15,800 rcf for 5 minutes at 4°C. The supernatant was analyzed by Liquid Chromatography (LC) - Tandem Mass Spectrometry (MS/MS). Calibration standards were prepared in blank plasma or brain homogenate by spiking 10 µL of standard solution and 10 µL of internal standard, followed by protein precipitation with 20 µL acetonitrile. The calibration range for lasmiditan was 1.1–1100 ng/mL in plasma and 0.5–550 ng/mL in brain. Chromatography used a C-18 column with a water/acetonitrile gradient. Detection was performed on a TSQ Quantum Ultra in positive ESI mode, monitoring transitions for lasmiditan (m/z 378→96, 378→159, 378→360) and aza-lasmiditan (m/z 379→159, 379→321, 379→335). The average brain and plasma concentration-time data were analyzed using PKSolver 2.0^>46^ with a non-compartmental approach.

Plasma and brain protein binding of Lasmiditan was assessed in WT and ASyn mice using rapid equilibrium dialysis (RED). A stock solution of lasmiditan (1.1 mg/mL) was prepared in DMSO and diluted to 40 µg/mL, then further diluted 200-fold in either mouse plasma or brain homogenate to a final concentration of 200 ng/mL, with a DMSO concentration of 0.5%. For the plasma study, 200 µL of plasma was added to the red chamber, and 350 µL of phosphate-buffered saline (PBS) to the white chamber. For the brain study, brain tissue was homogenized in twice the volume of PBS (e.g., 800 µL PBS for 400 mg brain), and 300 µL of this homogenate was placed in the red chamber, with 550 µL PBS in the buffer chamber. The RED devices were sealed and incubated at 37°C with shaking at 100 rpm for 4 hours (plasma) or 5 hours (brain). After dialysis, 50 µL samples were taken from both chambers, diluted with an equal volume of the opposing matrix, and analyzed by LC-MS/MS. The fraction of lasmiditan bound to proteins was calculated using peak area ratios of the analyte to the internal standard, following these equations: % free drug = 100 × (peak area ratio in buffer) / (peak area ratio in plasma or brain). % bound drug = 100 - % free drug.

### Behavioral Tests

#### Nest Building

Mice underwent a nest-building test as previously described^42–44^. About 1 hour before the dark phase, animals were placed in individual cages with all enrichment items removed. Nestlets (woven cotton pads for nesting) were weighed and provided to each cage (12–12.5 g per cage). Every 12 hours, an experimenter weighed the nestlet remaining in the feeder and any unused nestlet on the cage floor. The difference between the initial amount and the amount pulled down was recorded as ‘nestlet pulled down,’ while the difference between the amount pulled down but not shredded was recorded as ‘shredded nestlet’. Data for ‘nestlet pulled down’ and ‘shredded nestlet’ were collected over 72 hours at 12-hour intervals (12, 24, 36, 48, 60, and 72 h).

#### Y-Maze

Y-Maze testing was conducted as previously described^44^. Briefly, the mice were placed in the center of a Y-maze (three arms, each 30 cm long and 15 cm tall, arranged at 120-degree angles, and recorded from above. Entry into an arm was counted only when the mouse’s hind limbs completely crossed into it, with videos were analyzed for 7 minutes. Entries were grouped into triads, which were recorded when the mouse entered three different arms in a sequence starting from any arm. Trials were discarded if a mouse made fewer than 7 total entries (minimal exploration).

#### Open Field

Open field testing followed previously established protocols^42–44^. Animals were acclimated to the testing room for 2 hours over two days and tested 2 to 4 hours into their dark cycle. Each mouse was placed individually in the center of the open field arena (30 cm x 30 cm x 30 cm, with a 15 cm x 15 cm center grid marked on the floor) and allowed to explore undisturbed for 15 minutes, during which they were videotaped. After the testing, a blinded scorer analyzed the videos. Time spent moving, speed, and total distance traveled were recorded using ANY-Maze tracking software (Stoelting Co., Wood Dale, IL, USA).

#### Novel Object Recognition

As described before^44^, mice were individually placed in a 30 cm x 30 cm x 30 cm arena containing two identical objects (pineapple-shaped statues). They were allowed to explore for 10 minutes before returning to their home cage. After 2 hours, the mice were reintroduced to the arena for a second trial, where one familiar object (pineapple-shaped statue) and one novel object (a lighthouse statue) were presented. The mice explored the arena and objects for 10 minutes while being videotaped. After testing, videos were analyzed by visual inspection, with exploration defined as spending more than 7 seconds with an object. Data were presented as time spent with the familiar vs novel object as well as a Discrimination Index (DI) calculated using the formula: (time spent with the novel object – time spent with the familiar object) / (time spent with the novel object + time spent with the familiar object).

#### Inverted Screen Test

Briefly, a wire mesh (1 mm diameter wire with 12 mm squares, surrounded by a wooden frame 4 cm deep) was used^47^. After 2 hours of habituation, the mouse was placed in the center of the wire mesh screen and allowed to grip the wire mesh. The screen was inverted so that the mouse’s head was facing downwards first. The screen was held at a height of 40-50 cm above a padded surface. The time until mouse fell off the screen was recorded. Each mouse underwent three trials, with a total of three rounds performed for all mice.

### Quantitative PCR

DNA was isolated from the striatum, substantia nigra (SN), cortex, and hippocampus tissues using the Qiagen DNeasy Blood and Tissue Kit (QIAGEN, CA, USA). A total of 5 ng of isolated DNA was used for the quantification of relative mitochondrial DNA (mtDNA) content. The D-loop of the non-coding region in mtDNA was measured and normalized to the nuclear-encoded apolipoprotein B (ApoB) gene^48,49^. Quantitative PCR (qPCR) was performed using SsoAdvanced Universal SYBR Green Supermix (Bio-Rad, CA, USA). Fold changes in mtDNA content were calculated using the ΔΔCt method. Primers used: D-loop forward: (5’-CCCAAGCATATAAGCTAGTA-3’), Reverse: (5’- ATATAAGTCATATTTTGGGAACTAC-3’); ApoB forward: (5’-CGTGGGCTCCAGCATTCTA-3’), Reverse: (5’-TCACCAGTCATTTCTGCCTTTG-3’).

### Immunoblotting

Proteins were extracted from the striatum, SN, cortex, and hippocampus using RIPA buffer (50 mM Tris–HCl, 150 mM NaCl, 0.1% SDS, 0.5% sodium deoxycholate, 1% Triton X-100, pH 7.4) supplemented with a protease inhibitor cocktail (1:100), 1 mM sodium fluoride, and 1 mM sodium orthovanadate (Sigma-Aldrich, St. Louis, MO, USA). Protein concentration was determined using a bicinchoninic acid (BCA) assay. A total of 10–15 μg of protein was separated by electrophoresis using 10% SDS-polyacrylamide gels, followed by transfer to nitrocellulose membranes (Bio-Rad, Hercules, CA, USA). Membranes were blocked in 5% milk in TBST (Tris-buffered saline with 0.1% Tween 20) and incubated overnight with primary antibodies at 4°C with gentle agitation. Subsequently, membranes were incubated with the appropriate horseradish peroxidase-conjugated secondary antibody and visualized using chemiluminescence (Thermo Scientific, MA, USA) on an Azure 500 imaging system (Azure Biosystems, CA, USA). Optical density was quantified using Image Studio Lite software (LI-COR Biosciences, NE, USA), and protein levels were normalized to GAPDH. Primary antibodies used included PGC-1α (1:1000, NOVUS, CO, USA), TFAM (1:2000, Abcam, MA, USA), ATPsB (1:2000; Abcam, MA, USA), NDUFS1 (1:1000; Abcam, MA, USA), NDUFB8 (1:1000; Abcam, MA, USA), CD68 (1:3500; Bio-Rad, CA, USA), Iba1 (1:1000; NOVUS, CO, USA), aSyn (1:1000; BD Biosciences, NJ, USA), pSyn (1:500; Cell Signaling Technology, MA, USA), and GAPDH (1:2000; Cell Signaling Technology, MA, USA). More details of the antibodies can be found in the **Supplementary Table 3**. A single membrane was used to detect various combinations of target proteins, including (NDUFS1, ATPsB, TFAM, and GAPDH), (aSyn, PGC-1α, and GAPDH), (pSyn and GAPDH), (NDUFB8 and GAPDH), or (CD68, Iba1, and GAPDH). Although not all these combinations were employed in every experiment, GAPDH was consistently included on the same membrane as a reference standard.

### Measurement of mitochondrial oxidative phosphorylation

Mitochondria were isolated from the striatum, SN, and cortex using differential centrifugation as previously described^50^ (**Fig. 4a**). Briefly, tissues were homogenized in isolation buffer (210 mM mannitol, 70 mM sucrose, 1 mM EGTA, 5 mM HEPES, 0.5% (w/v) fatty acid-free BSA, pH 7.4) and centrifuged at 800 × g for 10 minutes at 4°C to remove nuclei and debris. The supernatant was subsequently centrifuged at 8,000 × g for 10 minutes at 4°C to pellet the mitochondria. The mitochondrial pellet was washed with isolation buffer and resuspended in a minimal volume for further analysis. Protein concentration was determined using BCA assay. The oxygen consumption rate (OCR) was measured using a Seahorse XF24 Analyzer (Agilent Technologies, Santa Clara, CA, USA) according to the manufacturer’s instructions. Isolated mitochondria were seeded in Seahorse XF24 plates at a concentration of 50 μg of protein per well and incubated in mitochondrial assay buffer composed of 2.0 mM HEPES (pH 7.2), 1.0 mM EGTA (pH 7.2), 70 mM sucrose, 220 mM mannitol, 10 mM KH₂PO₄, 5 mM MgCl₂, 0.2% (w/v) fatty acid-free BSA, 5 mM potassium pyruvate, 2.5 mM potassium malate, and 10 mM potassium glutamate. OCR was measured under basal conditions and after sequential injections of 4 mM ADP (to stimulate ATP production), 2 μM oligomycin (to assess ATP-linked respiration), 2 μM FCCP (as an uncoupler), and 2 μM rotenone/antimycin A (**Fig. 4b**). OCR values were normalized to mitochondrial protein content and expressed as pmol/min/μg protein. Data analysis was performed using Seahorse Wave software (Agilent Technologies).

### Immunohistochemistry

Following standard protocols, tissue sections were blocked using a solution containing 10% Normal Goat Serum, 0.5% Triton X-100, and 0.1 M Glycine in Tris Buffered Saline (TBS, pH 7.4) for 1 hour at room temperature (RT)^43,44^. Sections were then incubated overnight at RT with primary antibodies, including anti-phospho-alpha-synuclein D1R1R (1:500; Cell Signaling Technology, Danvers, MA, USA), anti-Iba1 (1:400; Fujifilm Irvine Scientific, Santa Ana, CA, USA), and anti-CD68 (1:200; Bio-Rad, Hercules, CA, USA). Primary antibodies were detected with secondary antibodies conjugated to Alexa Fluor 488 or 594 fluorochromes (Life Technologies-Molecular Probes, Grand Island, NY, USA) during a 2-hour incubation at RT. Sections were subsequently counterstained with 4’,6-diamidino-2-phenylindole dihydrochloride (DAPI; Life Technologies). Control conditions included the omission of primary or secondary antibodies, and isotype-specific antibodies or sera were used as substitutes for the omitted antibodies. More details of the antibodies used can be found in the **Supplementary Table 4**.

### Microscopy and Image Analysis

For the analysis of phosphorylated alpha-synuclein (pSyn), images were acquired using a Zeiss Axio Imager M2 (Zeiss, Oberkochen, Germany) or a Leica DMI6000 microscope (Leica Microsystems, Deerfield, IL, USA). In the hippocampus, three sections per animal were imaged, capturing distinct regions including the dentate gyrus, CA1, and CA3 at 40x magnification. For cortical regions, four sections per animal were imaged across three distinct areas at 20x magnification. Image analysis was performed using Fiji (NIH). Manual threshold adjustments were applied to each image to measure the percent area above the threshold, while total intensity measurements were obtained prior to thresholding. In the CA1 region of the hippocampus, the number of pSyn-positive cells was manually counted per frame. For Iba1 and CD68 analyses, images were captured using a Leica DMI6000 microscope with a 20x objective. Two tissue sections per animal were sampled in both the cortex and hippocampus. Within each cortical section, two regions were imaged, while three regions were captured within the hippocampus. Five-micron z-stacks were acquired, maximally projected using Leica software, and exported as individual channel TIF files. The total number of microglia was manually counted within each frame using the Iba1 channel. CD68 channel images were first analyzed for total intensity without preprocessing. Background subtraction was performed using the subtract function in FIJI ImageJ2 software (v2.14.0/1.54f). Subsequently, batch processing and thresholding were applied using default settings, and the percent area of staining was measured.

### Statistical analysis

GraphPad Prism 9 software (San Diego, CA, USA) was used for statistical analyses. For comparisons between three or more groups, one-way analysis of variance (ANOVA) followed by Tukey’s or Šídák’s post-hoc test for multiple comparisons between treatment groups was conducted. For analyzing the nest building task, a two-way repeated measures ANOVA followed by a Tukey’s post hoc test was used. Differences were considered significant at p < 0.05. Results are reported as mean ± SEM. Statistical details pertinent to each experiment are provided within the relevant result and legend sections.

## RESULTS

### Pharmacokinetics of lasmiditan in plasma and brain tissues

We examined the pharmacokinetic properties of lasmiditan in both plasma and brain tissues through protein binding analysis and Liquid Chromatography (LC) - Tandem Mass Spectrometry (MS/MS). Protein binding studies revealed 62±2.9% bound lasmiditan in the plasma of WT and 66±0.8% bound lasmiditan in the ASyn mice (**Supplementary Table 1**). In brain tissues, 83±1.0% bound Lasmiditan was measured in WT and 84±3.7% in the ASyn mice. For the LC-MS/MS measurements, lasmiditan was administered i.p. at 0.3mg/kg, 1 mg/kg or 2 mg/kg doses to WT and ASyn mice and plasma and brain samples collected at multiple timepoints. As shown in **Figure 1a-d** and **Supplementary Table 2**, the concentrations of total and free lasmiditan in plasma exhibited a dose-dependent increase in WT and ASyn mice. At all three doses, both free and bound lasmiditan concentrations were observed to peak at 0.5 hrs after i.p. dosing and decline gradually thereafter as a function of time. Levels were below the limits of detection at 24 hrs. In terms of maximum concentration (Cmax), both the WT and ASyn mice showed comparable values (**Supplementary Table 2**). For instance, at the 1.0 mg/kg dose, the Cmax for total lasmiditan was 129.8 ng/ml in WT and 106.1 ng/ml in ASyn and the Cmax for free lasmiditan was 48.03 ng/ml in WT and 36.07 ng/ml in ASyn. Similar Cmax trends were noted in the brain (**Fig. 1e-h and Supplementary Table 2**). Specifically, the 1.0 mg/kg dose resulted in a total lasmiditan Cmax of 295.01 ng/ml in WT and 252.93 ng/ml in ASyn and 44.32 ng/ml in WT and 36.26 ng/ml for free lasmiditan. It was also generally seen, in both brain and plasma, that although the increase in Cmax from 0.3 mg/kg to 1 mg/kg was more than threefold as expected, the change from 1.0 mg/kg to 2.0 mg/kg was less than twofold, indicating a deviation from linear pharmacokinetics (**Supplementary Table 2**).

In the brain, the time to reach maximum concentration (Tmax) for free lasmiditan remained constant across doses, with values ∼0.17 - 0.22 hours and the elimination half-life (t1/2) of 0.7 -1.04 hours across all doses and genotypes. Calculation of brain-to-plasma ratios indicated robust brain penetration with no notable differences across doses, genotypes, or conditions (**Fig. 1i**). For total lasmiditan, the brain-to-plasma ratios ranged from 2.27 to 2.46 in WT mice and 2.17 to 2.38 in ASyn mice. For free lasmiditan, the ratios ranged from 0.92 to 1.19 in WT and 0.98 to 1.24 in ASyn. These findings collectively indicate that lasmiditan exhibits consistent pharmacokinetic properties in both plasma and brain tissues, and good brain penetration, with no marked differences in transport efficiency or protein binding between WT and ASyn mice.

### Lasmiditan improves cognitive function in the young ASyn mice

We next assessed the effects of lasmiditan on the behavior and pathology of young (4-5.5 months of age) and old (10-11.5 months of age) ASyn mice, in the context of early and more advanced PD (**Fig. 2A**). Given the pharmacokinetic profile discussed above and previous studies ^33,51^, the young mice were treated with 1 mg/kg lasmiditan (i.p., every other day, for 6 weeks) and assessed via downstream behavioral, cellular and molecular analyses. Behaviorally the mice were subjected to different tests which assessed cognitive and motor functions including Y-maze, and novel object recognition (NOR), nest building,and open field tasks. In the Y-maze task, a measure of working memory, WT mice demonstrated 59.66 ± 1.096 % alternations (**Fig. 2a**). In contrast, ASyn mice exhibited significantly reduced alternations (52.22 ± 1.362%) compared to WT mice (F_2,66_ = 9.259, p = 0.0002, one-way ANOVA with Tukey’s multiple comparisons test). Lasmiditan treatment improved alternation rates in ASyn-Las mice to 57.45 ± 1.227%, which was significantly higher than ASyn mice (p = 0.014) and comparable to WT mice (p = 0.3808). The number of entries into the arms of the Y-maze did not differ significantly among groups, indicating similar activity levels (**Fig. 2b**). In the NOR test, WT mice had an average positive discrimination index (DI) of 0.1057 ± 0.03529 (**Fig. 2c**). ASyn mice, on the other hand, showed a negative DI of -0.05537 ± 0.0323 (F_2,61_ = 5.549, p = 0.0082 compared to WT, One-way ANOVA with Tukey’s multiple comparisons test), reflecting impaired object recognition. Lasmiditan treatment (ASyn-Las) reversed the DI to WT levels (0.1046 ± 0.04685), a value significantly higher than that of untreated ASyn mice (p = 0.0256). Moreover, when the exploration of novel versus familiar objects was analyzed, distinct group differences were revealed (**Fig. 2d**). WT and ASyn-Las groups preferentially explored the novel object, with WT mice showing a mean exploration time of 56.18% for the novel object (N) compared to 43.82% for the familiar object (F) (F_5,118_ = 13.32, p < 0.0001 for N vs F, One-way ANOVA with Tukey’s multiple comparisons test), and ASyn-Las mice spending 57.1% of their time on the novel object compared to 42.9% on the familiar object (p < 0.0001 for N vs F). In contrast, ASyn mice did not explore the novel object more (F_5,118_ = 13.32, p = 0.2391 for N vs F, One-way ANOVA with Tukey’s multiple comparisons test).

In the nest building task, the amount of woven cotton pads (nestlets) pulled down from a bundle placed at the top of the cage was measured every 12 hours as an indicator of gross motor function and motivation (**Fig. 2e**). WT mice successfully pulled down almost all the cotton within 72 hours, with a final mean value of 94.38 ± 3.812. In contrast, ASyn mice pulled down significantly less cotton compared to WT mice by the end of the test (34.11 ± 5.458, p < 0.0001, Two-way ANOVA with Tukey’s multiple comparisons test). Lasmiditan treatment did not improve this deficit, with the ASyn-Las group also showing significantly lower values than WT mice (26.36 ± 4.152, p < 0.0001). No significant difference was observed between the ASyn-Las and untreated ASyn groups at any time point (p > 0.05, Two-way RM-ANOVA with Tukey’s multiple comparisons test). The amount of shredded cotton in the cage, a measure of fine motor function was also assessed (**Fig. 2f)**. WT mice gradually shredded most of the cotton over time, with a final mean value of 55.1 ± 4.152 at 72 hours. In contrast, ASyn mice exhibited low shredding activity, compared to the WT mice with a final mean of 6.288 ± 3.812 (p < 0.0001, Two-way RM-ANOVA with Tukey’s multiple comparisons test). Lasmiditan treatment did not enhance the shredding activity of ASyn mice (7.4 ± 4.152 at 72 hours, p > 0.05 compared to ASyn).

With respect to the open field test, the total distance traveled differed significantly among the groups (**Fig. 2g)**. WT mice traveled a mean distance of 31.93 ± 2.201, while ASyn mice traveled slightly farther at 39.11 ± 3.182, although this difference was not statistically significant (F_2,71_ = 4.419, p = 0.2166, One-way ANOVA with Šídák’s multiple comparisons test). ASyn-Las traveled the greatest distance, with a mean of 43.22 ± 3.234, which was significantly greater than that of WT mice (p = 0.0141, One-way ANOVA with Šídák’s multiple comparisons test) but not significantly different from untreated ASyn mice (p = 0.6997, One-way ANOVA with Šídák’s multiple comparisons test). In terms of time spent in the central region of the field (vs the walls), a measure of anxiety-like behavior, lasmiditan treatment significantly increased the mean center time compared to untreated ASyn mice (218.3 ± 26.84 versus 144.9 ± 18.22 seconds, F_2,65_ = 3.7, p = 0.048, One-way ANOVA with Šídák’s multiple comparisons test) (**Fig. 2h**). In fact, ASyn-Las mice spent similar amounts of time in the central area akin to WT mice (213.3 ± 14.56 seconds, p = 0.8632, F_2,65_ = 3.7, One way ANOVA with Šídák’s multiple comparisons test). Overall, these data indicated an improvement in cognitive and affective functions but not motor performance upon lasmiditan treatment.

### Lasmiditan enhances mitochondrial biogenesis in the young ASyn mice

To assess the cellular and molecular effects of lasmiditan, associated with its behavioral effects, downstream assessments were performed in four brain regions: striatum, substantia nigra (SN), hippocampus, and cortex. Since most effects were found to be limited to the striatum, hippocampus and cortex, data from the SN have been presented in the text but not in the figures.

First, we assessed mitochondrial biogenesis (MB) by examining mitochondrial DNA replication as well as the expression of different mitochondrial proteins using qPCR and western blotting, respectively (**Fig. 3**). In the striatum, lasmiditan treatment significantly amplified relative mtDNA levels, compared to sham treated ASyn mice (F_2,21_ = 3.973, p = 0.0267, One-way ANOVA with Tukey’s multiple comparisons test) (**Fig. 3a**). mtDNA levels were also found to be significantly higher in the lasmiditan treated animals, compared to WT mice in the hippocampus (F_2,18_ = 10.17, p = 0.0009, One-way ANOVA with Tukey’s multiple comparisons test) (**Fig. 3b**). However, there was no significant difference between ASyn and ASyn-Las mice (p = 0.2755). No significant differences between any of the groups were found in the cortex (**Fig. 3c**). In the SN, significantly higher mtDNA was detected in the ASyn-Las mice (1.36 ± 0.18) than ASyn animals (0.79 ± 0.05) (F_2,21_ = 4.232, p = 0.0237, One-way ANOVA with Tukey’s multiple comparisons test).

**Figure 3.**
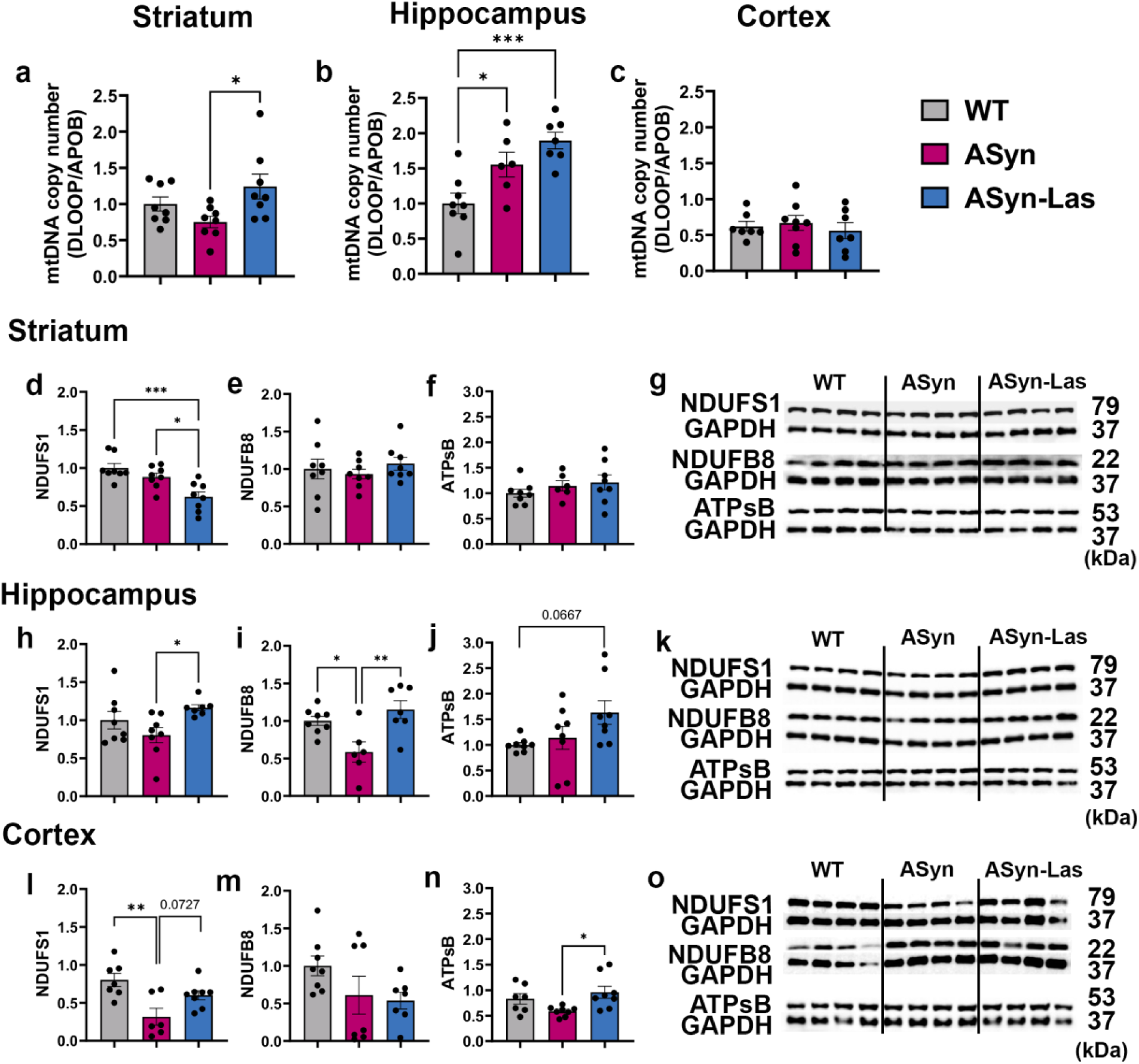
Mitochondrial DNA replication and mitochondrial protein expression in young mice. Relative mitochondrial DNA (mtDNA) levels were quantified by qPCR in the striatum (a), hippocampus (b), and cortex (c). Values were normalized to the mean mtDNA level of WT mice in each region. *** p = 0.001, * p < 0.05; one-way ANOVA with Tukey’s multiple comparisons test; mean ± SEM, n = 6–8/group. The expression levels of MB-relevant proteins NDUFS1, NDUFB8, and ATP synthase beta (ATPsB) were analyzed in the striatum, hippocampus, and cortex of the same groups of mice through western blotting. Striatal expression of NDUFS1 (d), NDUFB8 (e), and ATPsB (f) is shown, with representative blot images in (g). Similarly, hippocampal expression of the three proteins is shown in (h-j) with blot images in (k). Cortical expression data of NDUFS1, NDUFB8, and ATP synthase beta (ATPsB) is in (l-o). Protein levels were quantified relative to GAPDH and normalized to the mean expression level of WT mice in each region. **** p < 0.0001, ** p < 0.01, * p < 0.05; One-way ANOVA with Tukey’s multiple comparisons test; mean ± SEM, n = 8/group.

**Figure 4.**
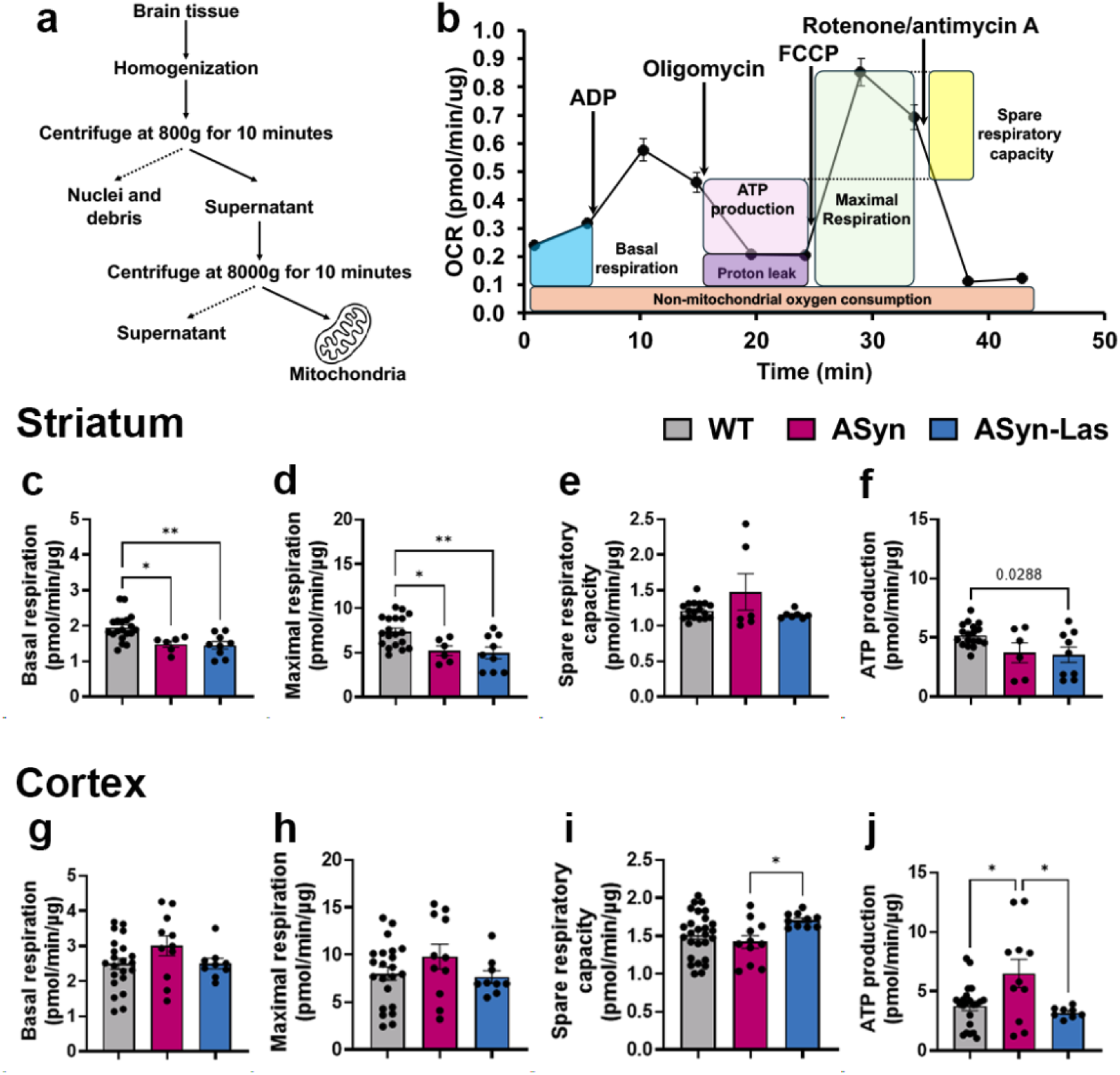
Assessment of mitochondrial function in young mice. (a) A schematic diagram illustrating the process of isolating mitochondria from brain tissues. (b) Oxygen consumption rate (OCR) profiles measured over 10 time points during the Mito Stress test using the Seahorse XF24 Analyzer. The graph shows OCR (pmol/min/µg protein) over time (min) in response to sequential injections of ADP, oligomycin, FCCP, and rotenone/antimycin A. Mitochondrial function in the striatum is represented in panels (c–f), and in the cortex in panels (g–j). Basal respiration is shown in (c) and (g), maximal respiration in (d) and (h), spare respiratory capacity in (e) and (i), and ATP production in (f) and (j). All values are presented as OCR (pmol/min/µg protein). Spare respiratory capacity is calculated as the difference between the maximal respiration after FCCP injection and basal respiration: Spare Respiratory Capacity = (Max rate after FCCP) – (Basal rate). ** p < 0.01, * p < 0.05; one-way ANOVA with Tukey’s multiple comparisons test; mean ± SEM, n = 6–27 individual wells/group.

With respect to mitochondrial protein expression, in the striatum, lasmiditan treatment did not improve the levels of either of the three markers assessed, namely NDUFS1, NDUFB8 and ATP Synthase Beta (ATPsB) (**Fig. 3d-f**, representative blot images in **Fig. 3g**). However, in the hippocampus, ASyn-Las mice showed significantly increased NDUFS1 expression compared to ASyn mice (F_2,20_ = 3.564, p = 0.0384, one-way ANOVA with Tukey’s multiple comparisons test) (**Fig. 3h**, representative blot images in **k)**. Hippocampal NDUFB8 also showed higher levels in the ASyn-Las mice compared to the sham treated ASyn animals (F_2,18_ = 7.391, p = 0.0041, One-way ANOVA with Tukey’s multiple comparisons test) (**Fig. 3i**, blot images in **k)**. NDUFB8 expression was also significantly lower in ASyn mice than WT (F_2,18_ = 7.391, p = 0.0289, One-way ANOVA with Tukey’s multiple comparisons test), while no significant differences were observed between WT and ASyn-Las mice (p = 0.5425). No differences between the groups were observed with respect to ATPsB in this region although ASyn-Las mice showed a strong increasing trend (p = 0.066 vs WT) (**Fig. 3j**, blot images in **k**). In the cortex, NDUFS1 expression was significantly reduced in ASyn mice compared to WT mice (F_2,18_ = 7.735, p = 0.0027, One-way ANOVA with Tukey’s multiple comparisons test) (**Fig. 3l**, representative blot images in **o**). Importantly, the ASyn-Las mice showed a almost statistically significant elevation (p = 0.0727), when compared to the ASyn group and was also not statistically different from WT (p = 0.2115). Cortical ATPsB expression was seen to be significantly higher in ASyn-Las mice compared to ASyn mice (F_2,20_ = 4.470, p = 0.0208, One-way ANOVA with Tukey’s multiple comparisons test) and no group differences were noted with respect to NDUFB8 (**Fig 3m-n**). Finally, no significant increases in either of the three proteins were noted in the SN upon lasmiditan treatment. Overall, these data indicated an upregulation of mitochondrial DNA and proteins, particularly in the hippocampus and cortex.

### Mitochondrial function is improved by lasmiditan in the young ASyn mice

Live mitochondria were isolated from the brain tissues of the young mice and their oxidative phosphorylation function was measured using the Seahorse XF24 Analyzer (Agilent Technologies, Santa Clara, CA, USA) via a Mito stress test (**Fig. 4a and b**). In the striatum, basal respiration was significantly reduced in ASyn and ASyn-Las compared to WT (F₂,₃₁ = 8.356, p = 0.0013, One-way ANOVA with Tukey’s test; WT vs. ASyn: p = 0.0181; WT vs. ASyn-Las: p = 0.0033) (**Fig. 4c**), and there were no significant differences between ASyn and Asyn-Las groups. Maximal respiration showed a similar profile (F₂,₃₁ = 7.547, p = 0.0021, One-way ANOVA with Tukey’s test; WT vs. ASyn: p = 0.0312; WT vs. ASyn-Las: p = 0.0045) (**Fig. 4d**). ATP production and spare respiratory capacity did not show significant differences between any of the groups (**Fig. 4e, f**). In the cortex, both basal and maximal respiration did not show significant differences between any of the groups, although the ASyn-Las mice showed a tendency to return to basal respiration levels seen in WT (**Fig. 4g, h**). Spare respiratory capacity on the other hand was significantly increased in ASyn-Las compared to ASyn groups (F₂,₅₇ = 7.469, p = 0.0013; ASyn vs. ASyn-Las: p = 0.034) (**Fig. 4i**). Interestingly, ATP production was significantly higher in ASyn compared to WT and lasmiditan treatment reduced it back to WT levels (F₂,₃₉ = 5.474, p = 0.0080; WT vs. ASyn: p = 0.0116; ASyn vs. ASyn-Las: p = 0.023) (**Fig. 4j**). These results suggest that while striatal mitochondrial function was not affected by lasmiditan, it was significantly altered in the cortex.

### Lasmiditan reduces α-synuclein and phosphorylated α-synuclein in the cortices of young mice

The expression of α-synuclein (aSyn) and phosphorylated α-synuclein (pSyn) were analyzed in all four regions of the young mice through immunoblotting and immunohistochemistry (**Fig. 5**). With respect to the immunoblotting, in the striatum, as expected, aSyn (**Fig. 5a, d**) and pSyn (**Fig. 5e, h**) expression were significantly increased in ASyn mice compared to WT mice (aSyn: F_2,21_ = 96.14, p < 0.0001 One-way ANOVA with Tukey’s multiple comparisons test; pSyn: F_2,21_ = 63.74, p < 0.0001, One-way ANOVA with Tukey’s test). However, no significant differences were observed between ASyn and ASyn-Las mice (**Fig. 5a, d**). The hippocampus showed a similar trend, with increased aSyn and pSyn in the ASyn mice with no changes upon lasmiditan treatment (**Fig. 5b, f**, representative blot images in **d, h**). However, in the cortex, although aSyn expression was significantly elevated in ASyn mice compared to WT (F_2,18_ = 86.44, p < 0.0001, One-way ANOVA with Tukey’s multiple comparisons test), the ASyn-Las mice exhibited a significant reduction in aSyn expression (p = 0.0048, **Fig. 5c**, representative blot images in **d**). Lasmiditan treatment also led to a significant decrease in pSyn expression in the ASyn-Las mice compared than ASyn mice (F_2,20_ = 23.63, p = 0.0286, One-way ANOVA with Tukey’s multiple comparisons test), though pSyn levels in ASyn-Las mice remained significantly higher than in WT mice (**Fig. 5g**, blots in **h**). No significant reductions in aSyn or pSyn were seen in the SN with lasmiditan treatment in any of the regions. These findings reveal that lasmiditan is effective in reducing aSyn and pSyn expression especially in the cortex. Given the immunoblotting changes, pSyn expression in the cortex was further analyzed using fluorescence immunohistochemistry (**Fig. 5i–l)**. In ASyn mice, pSyn was widely distributed throughout the observation fields, while lasmiditan treatment led to a visible reduction in pSyn expression (**Fig. 5i, j**). NIH Image J-based quantitative analysis showed that pSyn intensity was significantly higher in ASyn mice compared to WT (F_2,137_ = 129.9, p < 0.0001, One-way ANOVA with Tukey’s multiple comparisons test) (**Fig. 5l**). No significant differences in intensity were observed between ASyn and ASyn-Las mice (p > 0.9999). Conversely, while the percentage area (% area) of pSyn expression was significantly increased in ASyn mice compared to WT (F_2,137_ = 91.54, p < 0.0001, one-way ANOVA with Tukey’s multiple comparisons test), it was significantly reduced in ASyn-Las mice compared to ASyn mice (p < 0.0001), though it remained higher than in WT mice (p = 0.0285) (**Fig. 5k**). These findings indicate that lasmiditan effectively reduces both aSyn and pSyn expression in the cortex, particularly the spatial spread (% area) of pSyn.

**Figure 5.**
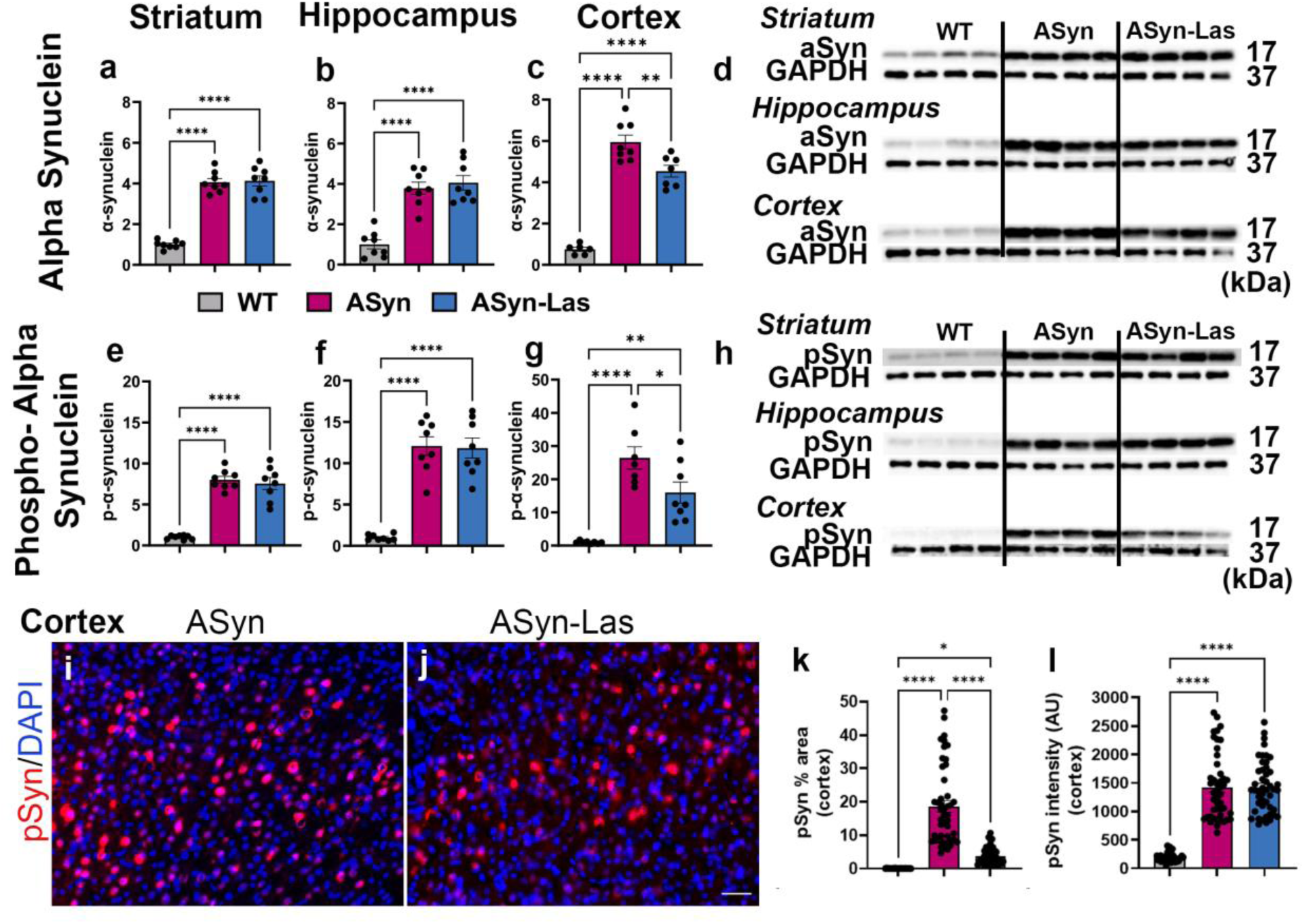
Expression of α-synuclein and phosphorylated α-synuclein in brain of young mice. The graphs show the expression levels of α-synuclein (aSyn) and phosphorylated α-synuclein (pSyn) in the striatum, hippocampus, and cortex by immunoblot quantification. The aSyn expression data is shown for the striatum (a), hippocampus (b), and cortex (c), with representative blot images in (d). The pSyn expression data is shown in (e-h). Quantification of aSyn and pSyn bands was performed relative to GAPDH, with all values normalized to the mean expression level of WT mice in each region, which was set to 1. Comparative immunohistochemical assessment of pSyn expression in the cortex in ASyn and ASyn-Las mice is shown via representative images in i and j. Quantitative analysis of pSyn intensity is shown in panel (l), while the percentage of pSyn-positive area (% area) is displayed in panel (k). **** p < 0.0001, ** p < 0.01, * p < 0.05; One-way ANOVA with Tukey’s multiple comparisons test. Data are presented as mean ± SEM, with n = 8/group for immunoblotting analysis and n = 6/group for IHC analysis. Scale bars: i, j = 50μm

### Lasmiditan treatment does not affect microglial number and activation in the younger ASyn mice

Microglial activation is known to be linked with α-synuclein pathology ^44,52^. Thus, given the increased pSyn in the cortex, we investigated the features of cortical microglia using western blotting and immunohistochemistry (**Fig. 6**). Immunostaining for the pan microglial marker Iba1 (**Fig. 6a,b**), and double -labeling of CD68 (marker of activated microglia) with Iba1 (**Fig. 6d, e**), demonstrated no appreciable qualitative differences in the expression of these markers between ASyn and ASyn-Las mice. In support, quantification of microglial numbers showed no significant differences among WT, ASyn, and ASyn-Las groups (F₂,₁₈ = 1.518, p = 0.2458, One-way ANOVA with Tukey’s test) (**Fig. 6c**). However, the % area of CD68-positive regions was significantly increased in ASyn mice compared to WT mice (F₂,₁₈ = 9.797, p = 0.0013, One-way ANOVA with Tukey’s test)(**Fig. 6f)**. However, lasmiditan treatment did not cause any significant alteration in % area compared to the ASyn mice (p = 0.9878) (**Fig. 6f**). These findings suggest that while ASyn pathology is associated with increased microglial activation as indicated by % area, lasmiditan does not impact microglial numbers or their activation state in the cortex. Immunoblot analysis supported the IHC data and did not show any significant alterations in Iba1 or CD68 expression upon lasmiditan treatment (**Fig 6g-i**). We also assessed hippocampal regions and no significant impact of lasmiditan were observed there via immunohistochemistry or immunoblotting. Immunohistochemistry revealed no significant differences in the total number of microglia among WT, ASyn, and ASyn-Las groups (F₂,₁₉ = 2.494, p = 0.1092) or in the CD68 % area (F₂,₁₉ = 1.600, p = 0.2279). Similarly, immunoblotting showed no significant differences in CD68 or Iba1 levels between the ASyn and ASyn-Las groups (CD68: F₂,₂₁ = 1.683, p = 0.2099; Iba1: F₂,₂₁ = 63.74, p < 0.0001).

**Figure 6.**
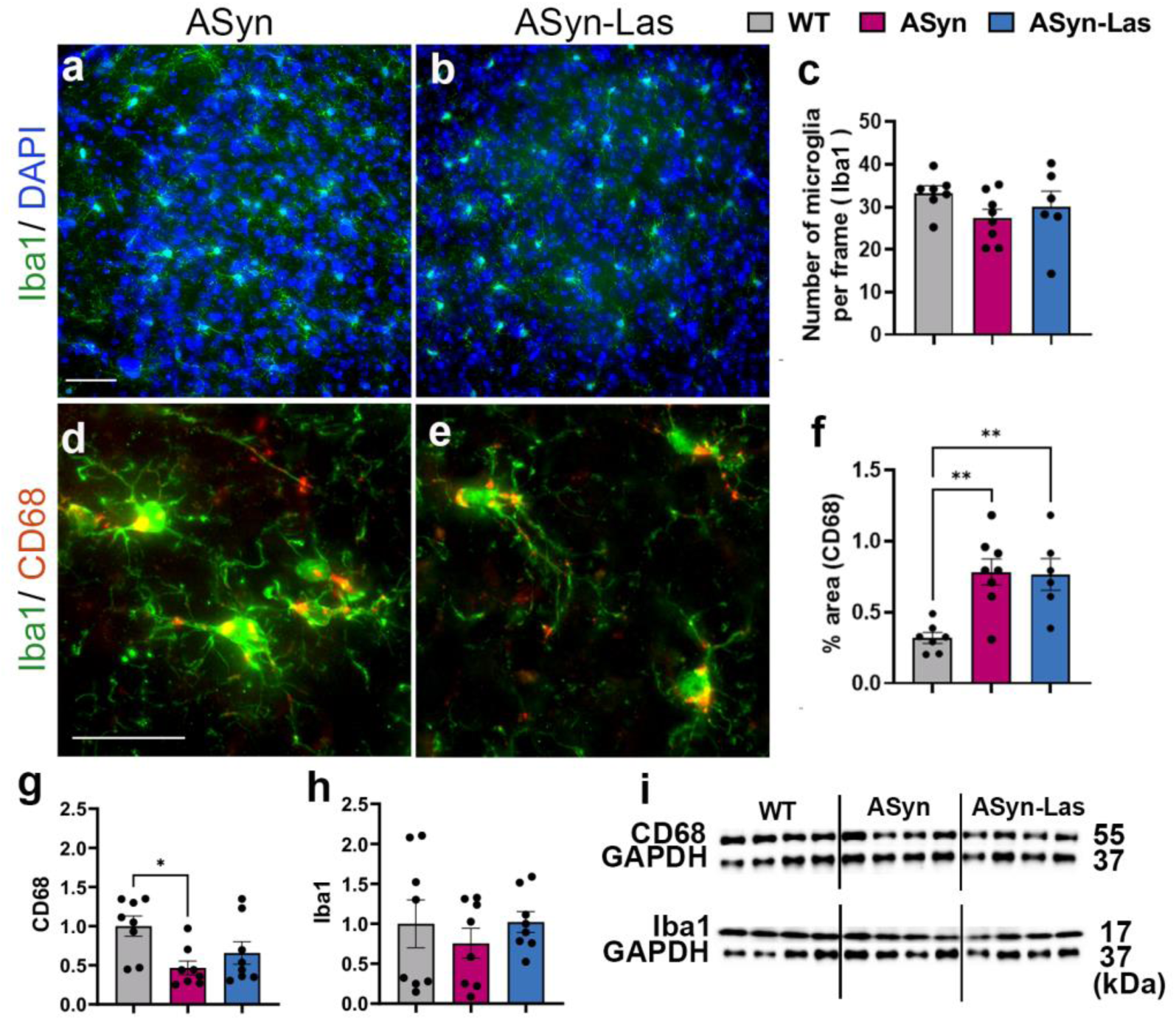
Effects of lasmiditan on microglial phenotype in young mice. The microglial expression of Iba1 and CD68 was analyzed using immunohistochemistry and immunoblotting in the cortex. Representative images of Iba1/DAPI and Iba1/CD68 immunostaining comparing ASyn and ASyn-Las mice is shown in (a, b, d and e). Quantification of the number of Iba1^+^ microglial cells per frame is shown in (c), and the percentage of CD68-positive microglial area (% area) is in (f). Total cortical protein levels of CD68 and Iba1 estimated via immunoblotting is shown in (g) and (h) with representative blot images displayed in (i). Quantification was performed relative to GAPDH, with all values normalized to the mean expression level of WT mice. ** p < 0.01, * p < 0.05; One-way ANOVA with Tukey’s multiple comparisons test. Data are presented as mean ± SEM, n = 8/group for immunoblotting and n = 6/group for IHC analysis. Scale bars: a,b, e, f = 50μm.

### Lasmiditan treatment improves cognitive function in the aged mice

The effect of lasmiditan in aged mice exhibiting advanced motor and cognitive symptoms was evaluated using an inverted screen test, novel object recognition and open field activity. In this study, given the higher age and advanced pathology in the old mice, another cohort of animals which received daily lasmiditan (ASyn-Las (D)) was included in addition to the alternate day administration group (ASyn-Las). Thus, four groups were compared in all evaluations: WT, ASyn, ASyn mice administered lasmiditan every other day (ASyn-Las), and ASyn mice administered lasmiditan daily (ASyn-Las (D)).

In the inverted screen test (**Fig. 7a**), ASyn mice showed significantly lower mean latencies to fall of the inverted grid compared to WT mice (F_3,80_ = 27.56, p < 0.0001, One-way ANOVA, with Tukey’s test). No significant differences were seen between the ASyn, ASyn-Las, and ASyn-Las (D) groups (p > 0.05). There was also no motor improvement seen in the open field task. No significant differences were observed in the total distance traveled during the open field test among the ASyn, ASyn-Las, and ASyn-Las (D) groups (F₃,₇₀ = 2.045, p = 0.1154, One-way ANOVA with Tukey’s test; ASyn: 16.49 ± 2.32 cm, ASyn-Las: 17.02 ± 2.19 cm, ASyn-Las (D): 21.99 ± 2.99 cm). However, when the mice were subjected to cognitive NOR tests, the lasmiditan treated mice performed better and showed greater exploration of the novel object vs the ASyn mice (F_7,132_ = 9.232, p < 0.0001, One-way ANOVA) (**Fig. 7c)**. ASyn-Las mice explored the novel object more extensively (novel: 58.68 ± 3.474 vs. familiar: 41.32 ± 3.474, p = 0.0019), and ASyn-Las (D) mice also showed a tendency to explore the novel object more (novel: 57.34 ± 2.206 vs. familiar: 42.66 ± 2.206, p = 0.4553). WT mice spent significantly more time on the novel object compared to the familiar object (p < 0.0001), while the sham ASyn mice showed no preference (p > 0.05). These results translated into a higher mean DI, in the lasmiditan treated mice than the sham ASyn mice with no significant differences found among the four groups (F_3,70_ = 1.346, p = 0.26, One-way ANOVA with Tukey’s test) (**Fig. 7b**). These data suggest that lasmiditan exerts a small but positive effect on cognition.

**Figure 7.**
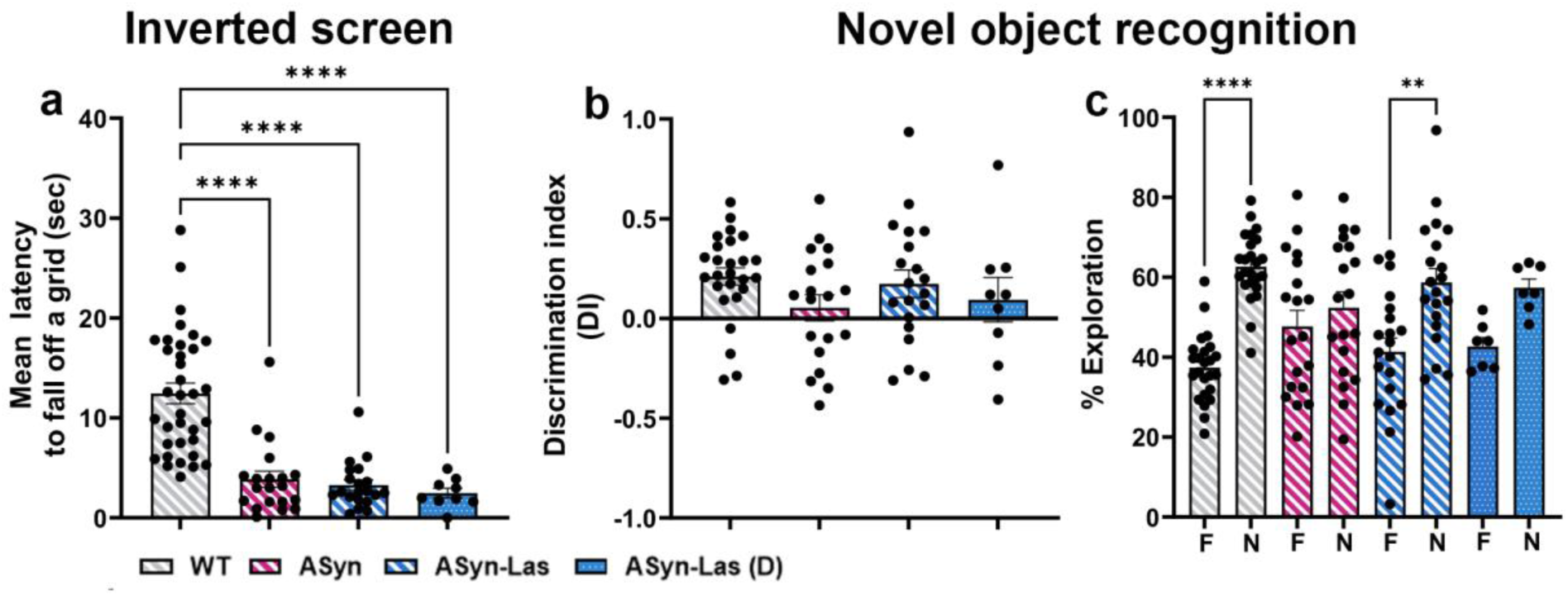
Behavioral assessment of aged mice. (a) shows data comparing the motor function in aged WT (gray bars with white diagonal stripes), ASyn (magenta bars with white diagonal stripes), ASyn-Las (blue bars with white diagonal stripes), and ASyn Las (D) (blue bars with white dots) mice via the Inverted screen test. Cognitive function was tested through an NOR test where the discrimination index (DI) (b) and novel vs familiar exploration (c) was measured. ** p < 0.01, * p < 0.0001; one-way ANOVA with Tukey’s multiple comparisons test. Data are presented as mean ± SEM, n = 9-35/group

### Mitochondrial biogenesis and mitochondrial function are minimally affected by lasmiditan in the aged mice

Assessment of mitochondrial DNA (mtDNA) replication showed no differences across all four groups in either the striatum or the hippocampus (**Fig. 8a, b**). In the cortex, the ASyn-Las (D) group had significantly higher mitochondrial DNA than the ASyn group (F_3,23_ = 3.508, p = 0.02, One-way ANOVA with Tukey’s test) and the WT group (p = 0.04) (**Fig. 8c**). The ASyn-Las mice on the other hand did not show differences from any of the other groups (p>0.05).

**Figure 8.**
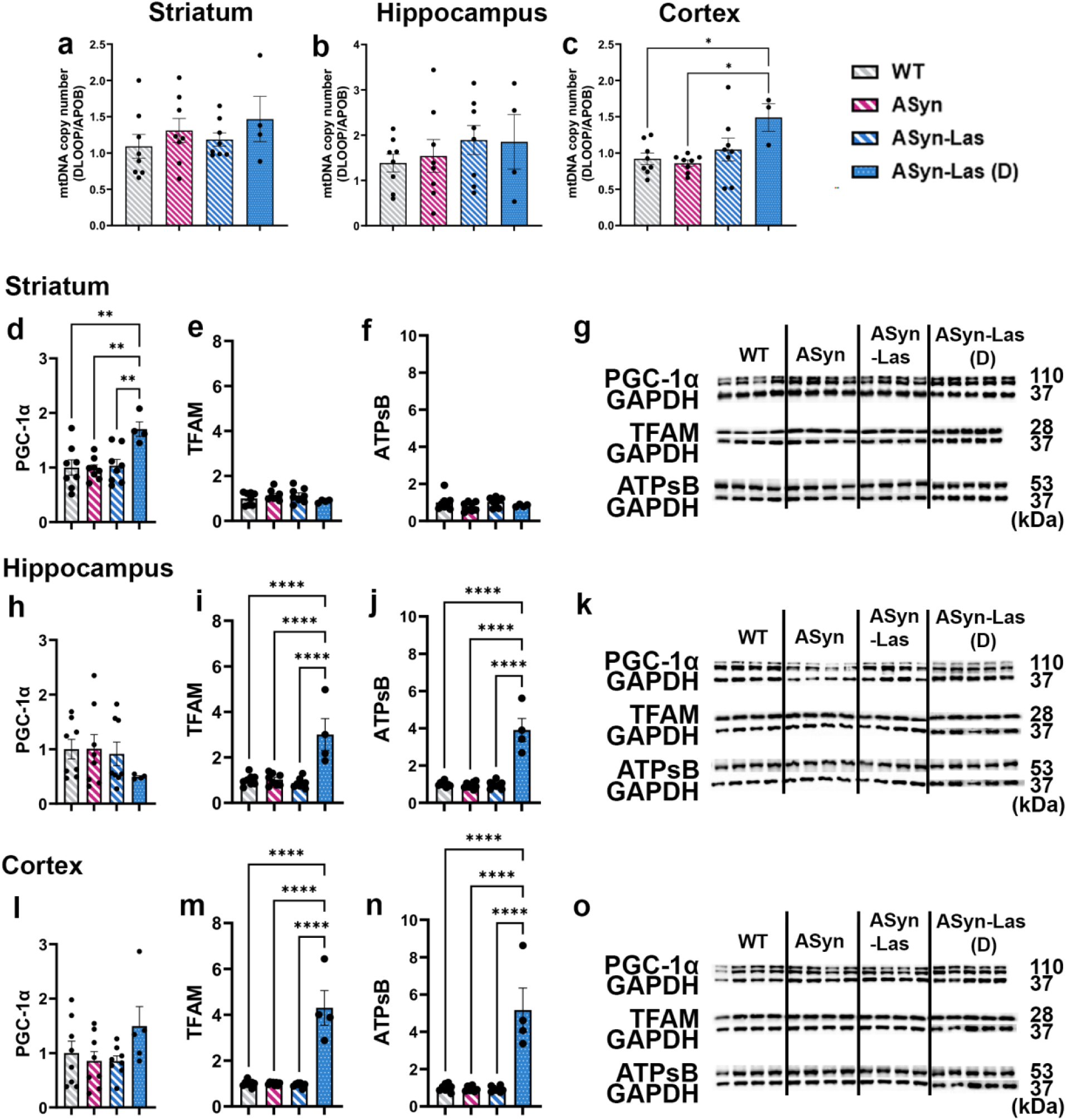
Induction of mitochondrial biogenesis in the aged mice. Mitochondrial DNA (mtDNA) levels were evaluated using qPCR, and the expression levels of MB-relevant proteins PGC-1α, TFAM, and ATPsB were analyzed using immunoblotting in the striatum, hippocampus, and cortex of old mice. (a-c) depict qPCR data from measuring mitochondrial DNA levels in the striatum (a), hippocampus (b), and cortex (c). All mtDNA values were normalized to the mean mtDNA level of WT mice in each region, which was set to 1. Estimation of striatal protein expression of PGC-1α, TFAM, and ATPsB via western blotting is shown in (d-f), with representative blot images in (g). Hippocampal and cortical expression of the three proteins is in (h-j) and (i-n) with representative blots shown in (k) and (o). Quantification of bands was performed relative to GAPDH, and all values were normalized to the mean expression level of WT mice in each region, which was set to 1. **** p < 0.0001, ** p < 0.01, * p < 0.05; One-way ANOVA with Tukey’s multiple comparisons test. Data are presented as mean ± SEM. For mtDNA analysis, n = 8/group for WT, ASyn, and ASyn-Las, and n = 4/group for ASyn-Las (D). For western blotting analysis, n = 8/group.

In terms of mitochondrial proteins, mean PGC-1α expression was significantly increased in the striata of the ASyn-Las (D) group compared to WT, ASyn, and ASyn-Las mice (F_3,24_ = 5.756, One-way ANOVA with Tukey’s multiple comparisons test: ASyn-Las (D) vs. WT, p = 0.0063; ASyn-Las (D) vs. ASyn, p = 0.0046; ASyn-Las (D) vs. ASyn-Las, p = 0.0093) (**Fig. 8d**). No significant differences across the groups were seen in the hippocampus and cortex (**Fig 8h, l**). With respect to TFAM, no significant differences in expression were observed among the four groups in the striatum (F₃,₂₄ = 1.18, p = 0.3383, One-way ANOVA with Tukey’s multiple comparisons test) (**Fig. 8e**). However, in the hippocampus and cortex, TFAM expression was significantly increased in the ASyn-Las (D) group compared to the WT, ASyn, and ASyn-Las groups (hippocampus: F₃,₂₄ = 16.10, p < 0.0001; cortex: F₃,₂₄ = 42.73, p < 0.0001, One-way ANOVA with Tukey’s multiple comparisons test) (**Fig. 8i, m**). No differences in TFAM levels were observed between the alternate-day lasmiditan treatment group and the ASyn group in either region (p > 0.05). Significant increases in ATPsB expression were also seen in the daily lasmiditan-treated group in both the hippocampus (F₃,₂₄ = 45.42, p < 0.0001) (**Fig. 8j**) and cortex (F₃,₂₄ = 28.10, p < 0.0001) (**Fig. 8n**), while the alternate-day lasmiditan regimen did not produce any increases in either region (p > 0.05). Similarly, no significant differences in ATPsB levels were observed across the groups in the striatum (F₃,₂₄ = 1.379, p = 0.2731, One-way ANOVA with Tukey’s multiple comparisons test) (**Fig. 8f**). In all three regions, the expression levels of TFAM and ATPsB in ASyn mice were not significantly different from those in WT mice, suggesting that α-synuclein overexpression alone does not affect basal mitochondrial protein levels. Representative blots for PGC-1α, TFAM, and ATPsB are provided in **Fig. 8g, k and o**. In terms of mitochondrial function (measured on live mitochondria via a Seahorse MitoStress test), no significant differences between ASyn and ASyn-Las groups were observed in the striatum (**Supp. Fig. 1**) while higher spare respiratory capacity was measured in ASyn-Las mice compared to ASyn and WT mice (F₂,₅₇ = 7.469, p = 0.0013; WT vs. ASyn-Las: p = 0.0008; ASyn vs. ASyn-Las: p = 0.034).

### Lasmiditan treatment does not affect α-synuclein expression or microglial activity in the context of advanced PD

Immunoblotting showed that although aSyn levels were significantly higher in the ASyn mice than WT mice as expected, in the striatum, hippocampus and cortex, neither types of lasmiditan treatment reduced aSyn expression (**Fig. 9a-d**). Similarly, in terms of pSyn, lasmiditan treatment did not show an effect in the hippocampus and cortex (**Fig. 9f,g**). Nevertheless, daily lasmiditan successfully reduced striatal pSyn expression compared to ASyn mice (F_3,22_ = 10.66, p = 0.0193, One-way ANOVA with Tukey’s test) (**Fig. 9e, h**). Additional immunofluorescence analysis revealed that although overall pSyn expression was not changed in the hippocampus, distinct regional patterns, especially in the CA3 and dentate gyrus (DG) were noted (**Fig. 9i-n**). In the CA3 region, ASyn mice displayed significant increases in both pSyn intensity and % area compared to WT mice (**Fig 9i, o**). Lasmiditan treatment reduced pSyn intensity in both the alternate-day and daily treatment groups (F₃,₄₄ = 20.94, p < 0.0001, One-way ANOVA with Tukey’s test) (**Fig. 9j,k,o**). However, % area showed decreasing trends but not significant reductions in the lasmiditan-treated groups, despite a significant increase in ASyn mice compared to WT (p = 0.0075) (**Fig. 9p**). In the DG region, pSyn intensity was significantly reduced in the daily lasmiditan-treated group compared to untreated ASyn mice (p = 0.0068, F₃,₄₄ = 17.74, One-way ANOVA with Tukey’s test), while the alternate-day treatment group did not show a significant reduction (**Fig. 9l,m.n,q**). WT mice had significantly lower pSyn intensity than all other groups (p < 0.0001) (**Fig. 9q**). In terms of % area of pSyn expression in the DG, no significant differences were observed across lasmiditan treatment groups, although ASyn mice exhibited an increase compared to WT (p < 0.0001) (**Fig. 9r**). No such IHC-based regional differences were seen in the cortex.

**Figure 9.**
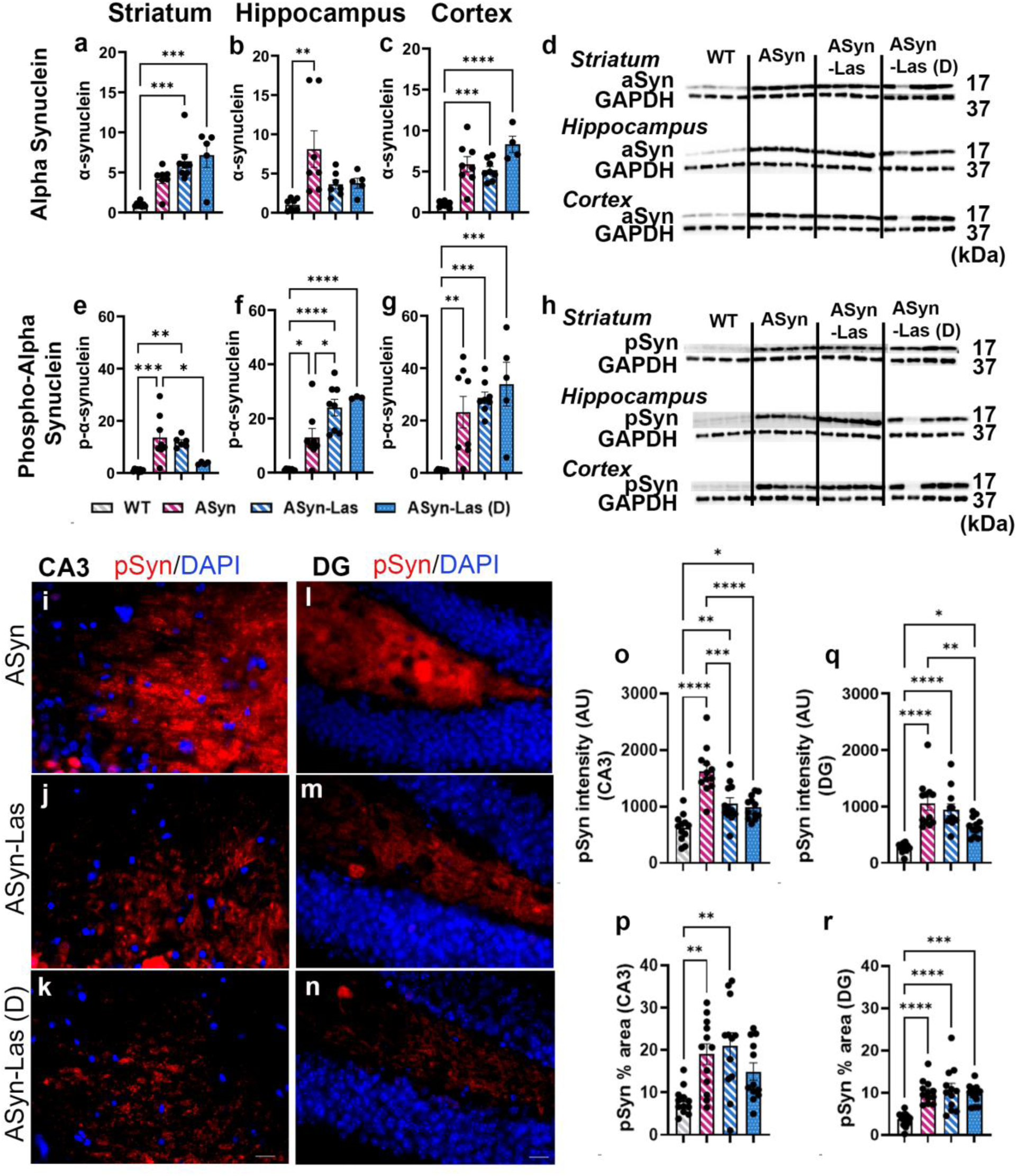
α-synuclein and phosphorylated α-synuclein expression in the aged mice. The expression levels of α-synuclein (aSyn) and phosphorylated α-synuclein (pSyn) were analyzed using immunoblotting and Immunohistochemical (IHC). Total aSyn protein expression in the striatum (a), hippocampus (b), and cortex (c), and pSyn in the striatum (e), hippocampus (f), and cortex (g) is shown. Representative immunoblot images are in (d) and (h). Quantification for immunoblotting was performed relative to GAPDH, and all values were normalized to the mean expression level of WT mice in each region, which was set to 1. IHC analysis of pSyn in the hippocampal CA3 and DG is shown in (i-r). Qualitative images of pSyn expression in the ASyn, ASyn-Las, and ASyn-Las (D) mice of the CA3 and DG are in (i-k) and (l-n) respectively. For the CA3, quantitative analysis of pSyn intensity is presented in panel (o), and the percentage of pSyn-positive area (% area) is displayed in panel (p). Similarly, pSyn intensity in the DG is quantified in panel (q), and the % area is displayed in panel (r). **** p < 0.0001, *** p < 0.001, ** p < 0.01, * p < 0.05; One-way ANOVA with Tukey’s multiple comparisons test. Data are presented as mean ± SEM, n = 8/group for immunoblotting and n = 6/group for pSyn IHC analysis. Scale bars: i-k = 20μm

Microglial activity in the hippocampus and cortex was examined (**Fig. 10)**. Across three regions of the hippocampus (CA1, CA3 and DG), the total number of Iba1-positive (Iba1^+^) microglia was significantly increased in ASyn mice compared to WT mice (35.47 ± 1.922 vs. 27.06 ± 0.9648, F₃,₂₁ = 8.941, p = 0.0358, One-way ANOVA with Tukey’s test) (**Fig. 10d**). However, lasmiditan treatment did not significantly alter microglial counts compared to the ASyn group (**Fig. 10a-c and d**). Regarding CD68 % area, no significant differences were observed among the four groups (F₃,₂₂ = 0.5278, p = 0.6678, one-way ANOVA with Tukey’s test) (**Fig. 10h, representative images are in e-g**). Moreover, no significant effects of lasmiditan were seen when the three regions were analyzed individually either. Immunoblot analysis for CD68, using tissues from the entire hippocampus, showed no significant differences between WT and ASyn groups, nor did alternate day lasmiditan treatment change CD68 expression compared to the sham ASyn group. However, daily lasmiditan treatment resulted in a notable reduction in CD68 expression (F₃,₂₄ = 3.697, one-way ANOVA with Tukey’s test; p = 0.066 for ASyn vs ASyn-Las (D) and p = 0.0256 for ASyn-Las vs ASyn-Las (D) (**Fig. 10i, k**). In terms of Iba1 levels, immunoblot analysis revealed no significant differences in expression across all four groups (WT, ASyn, ASyn-Las, ASyn-Las (D) (**Fig. 10j, k**). No changes in the cortex were observed upon lasmiditan treatment.

**Figure 10.**
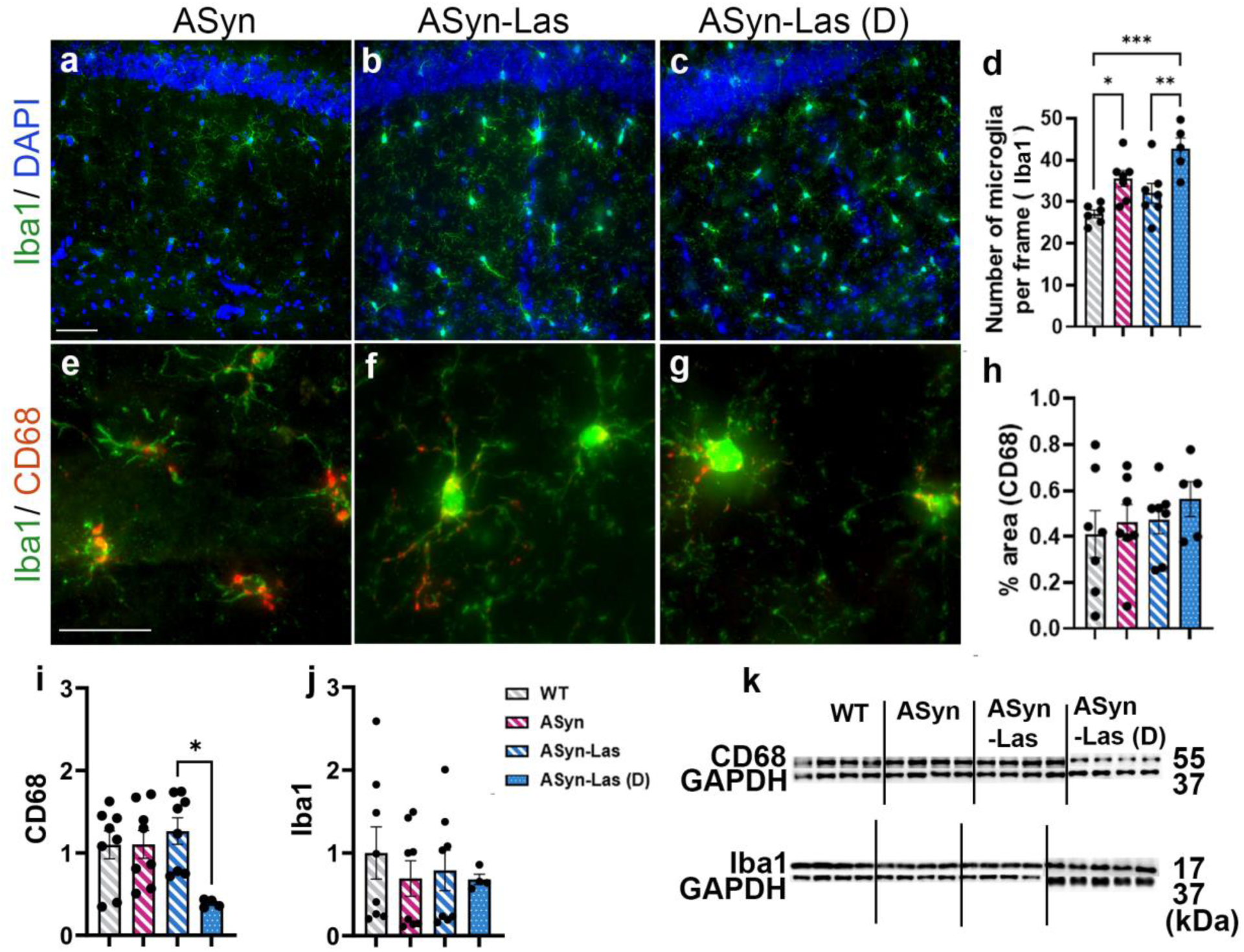
Characterization of lasmiditan’s effects on microglial activity in the aged mice. Panels (a-c) show representative images of Iba1/DAPI (blue) immunostaining for ASyn, ASyn-Las, and ASyn-Las (D) mice. Panels (e-g) display Iba1/CD68 co-staining. Associated quantification of the number of Iba1^+^ microglial cells per frame is shown in (d), and the percentage of CD68-positive microglial area (% area) is shown in (h). Western blotting based Iba1 and CD68 expression levels are depicted in (i) and (j) with representative blot images in (k). Quantification for immunoblotting was performed relative to GAPDH, with all values normalized to the mean expression level of WT mice. *** p < 0.001, ** p < 0.01, * p < 0.05; One-way ANOVA with Tukey’s multiple comparisons test. Data are presented as mean ± SEM, n = 8/group for immunoblotting and n = 6/group for IHC analysis. Scale Bars: 50μm.

## DISCUSSION

In essence, this study demonstrates that the selective 5-HT1F receptor agonist, lasmiditan, improves cortico-hippocampal pathology and mitigates PD-relevant cognitive deficits. Specifically, systemic lasmiditan treatment boosted MB and mitochondrial function, while lessening the α-synuclein load, especially in younger line 61 Thy1-aSyn mice. In PD, non-motor symptoms such as cognitive impairment significantly increase disease burden, beyond the classic motor symptoms^3–6^. Our findings suggest that targeting enhanced MB and α-synuclein reduction, through 5-HT1F receptor agonism, may represent an effective strategy to address cognitive symptoms in PD.

PK analysis showed that the peak plasma and brain levels of lasmiditan at the chosen 1mg/kg dose in WT mice were about 129 ng/ml and 295 ng/ml with a constant Tmax of around 0.5 hours across all three tested doses. Importantly, the brain-to-plasma concentration ratio was around 2, indicating good distribution into the brain. Moreover, PK characteristics were broadly similar between the WT and transgenic ASyn mice. In the context of human studies, total lasmiditan concentrations measured in mouse plasma were similar to what has been seen in human plasma^53^ (FDA, Clinical Pharmacology and Biopharmaceutics Review for NDA 211280: https://www.accessdata.fda.gov/drugsatfda_docs/nda/2019/211280Orig1s000ClinPharmR.pdf, accessed 1/9/2025). The protein binding of lasmiditan in mouse plasma was also comparable to that in humans; lasmiditan was noted to be ∼64% bound in mouse plasma and binding in human plasma is known to be around 55-60% (FDA, Printed Labeling for NDA 211280: https://www.accessdata.fda.gov/drugsatfda_docs/nda/2019/211280Orig1s000lbl.pdf). In brain homogenates, protein binding was ∼83%, which was higher than that in plasma, supporting the observed free brain-to-plasma concentration ratio being greater than 1. Additionally, a less than two-fold increase in Cmax and AUC between 1 and 2 mg/kg doses of lasmiditan was seen, suggesting a saturation effect. Interestingly, a similar, less than proportional increase in Cmax and AUC between 100 mg vs 200 mg dose is also noted in humans^54^.

In terms of behavioral effects, lasmiditan particularly improved cognitive function, in both the young and aged mice. In the younger (∼5.5 months old) ASyn mice, lasmiditan significantly enhanced the performance in both the Y-maze and NOR tasks. These tasks measure episodic and working memory, respectively, which are both aspects of cognition known to be affected early in PD. Thus, our data indicate that both these types of memory were improved by lasmiditan. The older (∼11.5 months old) mice were primarily tested using the NOR task. it was observed that the Asyn mice with lasmiditan were better able to distinguish the novel object from the familiar one, both in the everyday and every other day dosing regimens. To our knowledge, this is the first time that 5-HT1F agonism via lasmiditan has been shown to improve cognitive function in the context of PD. In contrast, lasmiditan did not alleviate motor dysfunction in either the young or old mice (nest building and open field activity). Such specificity of lasmiditan’s effect on cognitive function is interesting and warrants further investigation. However, a couple of factors can be envisioned to have contributed to this effect. Firstly, it is known that 5-HT1F receptor expression is especially high in the hippocampus and cortex, key regions mediating cognitive functions^34,55–57^. Secondly, the bioavailability of lasmiditan might have been higher in the hippocampus due to a more vulnerable blood brain barrier reported in this region^58,59^. Our data broadly shows good brain penetration by lasmiditan but does not address regional differences, which would be important to investigate in future studies.

At the molecular level, lasmiditan treatment significantly elevated the levels of MB-relevant proteins (NDUFS1, NDUFB8, and ATPsB) in the hippocampus and cortex while also increasing mtDNA content by approximately 10–15% in the hippocampus (**Fig. 3**). These changes were associated with changes in mitochondrial function in the cortex. Specifically, lasmiditan treatment ‘reset’ mitochondrial function, including basal respiration, maximal respiration, ATP production and spare respiratory capacity, to WT levels as opposed to compensatory increases in these parameters seen in the untreated ASyn controls. In the striatum, although mtDNA elevation was seen, this did not translate to alterations in mitochondrial function. In essence, these data signified improved mitochondrial biology in cognitively relevant regions, namely the hippocampus and cortex, thus supporting the observed behavioral improvements in episodic and working memory. Comparatively in the aged mice, although the alternate day administration of lasmiditan rendered no significant changes, daily lasmiditan treatment amplified mtDNA in the cortex as well as levels of MB-relevant proteins TFAM and ATPsB in both the hippocampus and cortex. A mild associated improvement in mitochondrial function in the cortex, in the form of increased spare respiratory capacity was noted, which may have ultimately bettered performance on the NOR task. This profile suggested a partial recovery of mitochondrial adaptability with age-related limitations. Fundamentally, the findings on mitochondrial function highlight the selective benefits of lasmiditan in cortical regions critical for cognition, while underscoring the challenges of reversing mitochondrial dysfunction in advanced PD.

With respect to α-synuclein, lasmiditan decreased cortical aSyn and pSyn protein levels by approximately 20% in the young mice while no significant reductions were seen in the hippocampus (**Fig. 5**). Immunohistochemistry further revealed that while pSyn intensity was not altered, the spatial distribution of pSyn (measured as % area) was reduced, indicating that lasmiditan had restricted the spread and expansion of pSyn aggregates in the cortex. In the older animals, no significant decreases in total aSyn or pSyn protein levels were seen in either the hippocampus or cortex even with daily lasmiditan treatment. However, careful immunohistochemical assessments in different hippocampal regions showed diminished pSyn intensity in the CA3 and DG areas upon daily lasmiditan administration. Thus, the pattern of pSyn distribution differed by age and brain region. Basically, while young cortical regions showed no change in pSyn intensity but a reduction in % area, the specific aged hippocampal regions exhibited reduced pSyn intensity without a significant decrease in % area. These differences suggest that the effects of interventions on pSyn pathology, whether focused on limiting spatial expansion or reducing density, depend on the disease stage and the specific brain region. In young cortical areas during early disease, the spread of aggregates, measured as % area, is effectively controlled. In contrast, in aged hippocampal regions, with more advanced pathology, interventions may reduce the density of aggregates (intensity) without significantly decreasing the affected area. These findings emphasize the complex and region-specific nature of the impact of MB induction on pSyn pathology.

Correlatively, no prominent effects on total Iba1 (pan microglial) or CD68 (activated microglial) protein expression, as well as immunoreactivity were seen upon lasmiditan treatment in the younger mice in the hippocampus or cortex. However, a trend towards returning microglial numbers to WT levels, particularly in the cortex was noted. The aged mice showed no significant changes in microglial numbers or CD68 expression, or notable trends, via immunohistochemistry; however, a significant reduction in total protein levels of CD68 was noted in the hippocampus with daily lasmiditan administration. Inflammation, particularly microglial activation, is known to be associated with α-synuclein pathology^52,60–62^. In this context, our data do not support a significant impact of lasmiditan treatment on microglial activity and suggest that the pSyn reducing effects of lasmiditan were probably independent of microglia. It is known that glial cells do not express 5-HT1F receptors^63^ which could have contributed to such an outcome. Nevertheless, any indirect effects of 5-HT1F agonism occurring through downstream neuron-glia cross talk cannot be completely ruled out.

In conclusion, our study indicates that MB induction by lasmiditan in young ASyn mice stabilizes the metabolic and synaptic environment of the cortico-hippocampal region and suppresses the spatial spread of pSyn aggregates, exerting a neuroprotective effect on cognition. While effects diminish in advanced disease, optimizing dosing regimens can partially restore MB-related molecular improvements. These findings emphasize the importance of timing and dosing in therapeutic interventions targeting mitochondrial dysfunction and synaptic instability. Lasmiditan, in the context of its FDA approved use for the acute treatment of migraine, exerts its therapeutic influence by suppressing cortical spreading depression (CSD) and excessive trigeminovascular responses^38,64–67^. CSD, characterized by a wave of neuronal and glial depolarization followed by suppression of activity, is associated with sensory circuit hyperactivation, vascular dysregulation, and the release of inflammatory mediators^38,68–70^. Although CSD itself is not a key feature of PD models, lasmiditan’s ability to modulate these pathways may stabilize sensory circuits and enhance mitochondrial and synaptic function in regions with high energy demands, such as the hippocampus and cortex. These regions, which are critical for cognition and memory, are particularly vulnerable to metabolic stress and synaptic dysfunction in PD^71–74^. Thus, repurposing lasmiditan may position it as a strong candidate to address PD’s non-motor symptoms as well as other neurological disorders characterized by metabolic and synaptic dysfunction.

## ACKNOWLEDGEMENTS

We thank Dr Jaroslav Janda for his technical help with respect to the immunoblotting, Dr Moulon Luo for his guidance with the Seahorse analyses, and Dr Ju Gao for his expert input on the mitochondrial isolations. The LC-MS/MS and protein binding work for PK analyses was conducted at the UA Analytical Chemistry Shared Resource (ACSR), and we thank Sherry Chow and Wade Chew in this regard. The UA ACSR is supported by Cancer Center Support Grant via the National Cancer Institute of the National Institutes of Health P30 CA023074 grant. Part of the microscopy was performed at the University of Arizona (UA) Imaging Core - UA RII Imaging Cores - Optical core facility, RRID:SCR_023355. The study was supported by a Michael J Fox Foundation grant (MJFF-021969) and UA Intramural funds to LM; NIH post-doctoral training grant (T32 AG044402) to AI, UA UBRP and Galileo Scholar program support for JM, and UA FRONTERA and MARC program support for PW.

## AUTHOR CONTRIBUTIONS

AI – Experimental design, Collection and assembly of data, Data analysis and Interpretation, Manuscript writing

JM – Collection and assembly of data, Data analysis

MJC – Collection and assembly of data, Data analysis, General technical support, Manuscript editing

KB – Collection and assembly of data, Data analysis and Interpretation, Manuscript editing

PW – Collection and assembly of behavioral data, Data analysis

NM – Analysis of behavioral data

PVS – Running and Data analysis support for qPCR experiments

RGS – Experimental design and conceptualization, Data interpretation and Resources support, and Manuscript editing

LM – Overall conception and design of the study, Data analysis and Interpretation, Manuscript writing, Financial support, Final approval of manuscript.

---

**Supplementary Table 1.** Protein binding of Lasmiditan in plasma and brain tissue.

|  | % Bound (mean±SD) | % Free (mean±SD) |
| --- | --- | --- |
| **Plasma** |  |  |
| Wild type | 62±2.9 | 37±2.9 |
| ASyn | 66±0.8 | 34±2.9 |
| **Brain** |  |  |
| Wild type | 83±1.0 | 17±1.0 |
| ASyn | 84±3.7 | 16±3.7 |

**Supplementary Table 2.** Pharmacokinetics of Lasmiditan in plasma and brain tissue.

|  | 0.3 mg/kg |  |  |  | 1.0 mg/kg |  |  |  | 2.0 mg/kg |  |  |  |
| --- | --- | --- | --- | --- | --- | --- | --- | --- | --- | --- | --- | --- |
|  | WT |  | ASyn |  | WT |  | ASyn |  | WT |  | ASyn |  |
|  | Plasma | Brain | Plasma | Brain | Plasma | Brain | Plasma | Brain | Plasma | Brain | Plasma | Brain |
| **Total Lasmiditan** |  |  |  |  |  |  |  |  |  |  |  |  |
| Cmax (ng/ml) | 28.31 | 69.74 | 12.82 | 28.99 | 129.8 | 295.01 | 106.1 | 252.93 | 179.8 | 415.94 | 135.5 | 294.04 |
| Tmax (h) | 0.5 | 1.23 | 0.5 | 1.13 | 0.5 | 1.14 | 0.5 | 1.19 | 0.5 | 1.16 | 0.5 | 1.09 |
| t1/2 (h) | 2.36 | 5.81 | 2.57 | 5.8 | 2.32 | 5.27 | 2.05 | 4.89 | 2.35 | 5.44 | 2.11 | 4.58 |
| AUC 0-t (ng/ml·h) | 86.1 | 212.09 | 38.8 | 87.73 | 332.4 | 755.47 | 293.4 | 699.43 | 431.74 | 998.76 | 378.99 | 822.41 |
| **Free Lasmiditan** |  |  |  |  |  |  |  |  |  |  |  |  |
| Cmax (ng/ml) | 10.47 | 12.5 | 4.36 | 5.41 | 48.03 | 44.32 | 36.07 | 36.26 | 66.53 | 69.65 | 46.07 | 44.93 |
| Tmax (h) | 0.19 | 0.22 | 0.17 | 0.21 | 0.19 | 0.17 | 0.17 | 0.17 | 0.19 | 0.19 | 0.17 | 0.17 |
| t1/2 (h) | 0.87 | 1.04 | 0.87 | 1.08 | 0.86 | 0.79 | 0.7 | 0.7 | 0.87 | 0.91 | 0.72 | 0.7 |
| AUC 0-t (ng/ml·h) | 31.86 | 38.01 | 13.19 | 16.38 | 122.99 | 113.5 | 99.76 | 100.27 | 159.74 | 167.25 | 128.86 | 125.68 |

WT: Wild type mice, ASyn: Thy1-aSyn mice, Cmax: Maximum concentration, Tmax: Time to maximum concentration, t1/2:
Elimination half-life, AUC<sub>0-t</sub>: Area under the concentration-time curve from time 0 to time t.

**Supplementary Table 3.**
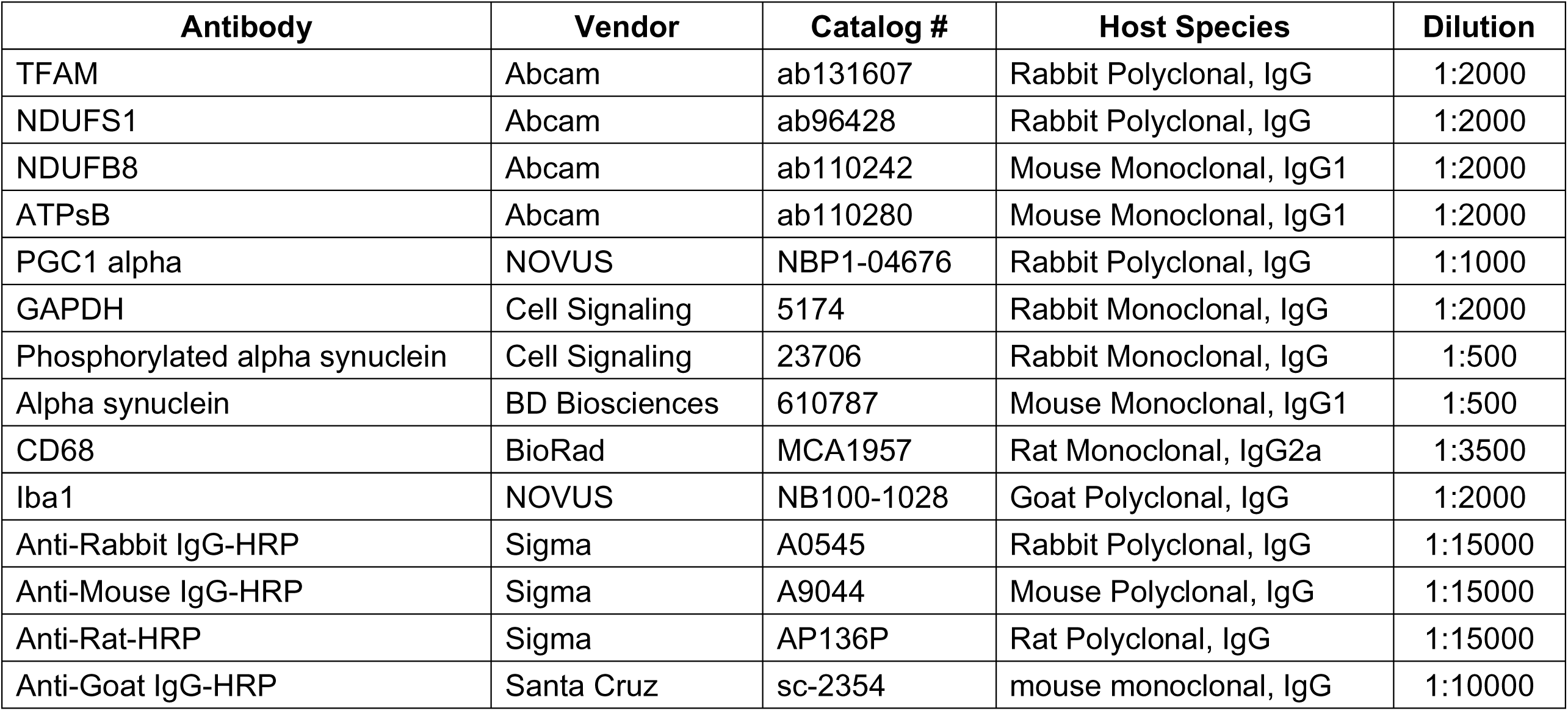
Antibody list for immunoblotting.

| **Antibody** | **Vendor** | **Catalog #** | **Host Species** | **Dilution** |
| --- | --- | --- | --- | --- |
| TFAM | Abcam | ab131607 | Rabbit Polyclonal, IgG | 1:2000 |
| NDUFS1 | Abcam | ab96428 | Rabbit Polyclonal, IgG | 1:2000 |
| NDUFB8 | Abcam | ab110242 | Mouse Monoclonal, IgG1 | 1:2000 |
| ATPsB | Abcam | ab110280 | Mouse Monoclonal, IgG1 | 1:2000 |
| PGC1 alpha | NOVUS | NBP1-04676 | Rabbit Polyclonal, IgG | 1:1000 |
| GAPDH | Cell Signaling | 5174 | Rabbit Monoclonal, IgG | 1:2000 |
| Phosphorylated alpha synuclein | Cell Signaling | 23706 | Rabbit Monoclonal, IgG | 1:500 |
| Alpha synuclein | BD Biosciences | 610787 | Mouse Monoclonal, IgG1 | 1:500 |
| CD68 | BioRad | MCA1957 | Rat Monoclonal, IgG2a | 1:3500 |
| Iba1 | NOVUS | NB100-1028 | Goat Polyclonal, IgG | 1:2000 |
| Anti-Rabbit IgG-HRP | Sigma | A0545 | Rabbit Polyclonal, IgG | 1:15000 |
| Anti-Mouse IgG-HRP | Sigma | A9044 | Mouse Polyclonal, IgG | 1:15000 |
| Anti-Rat-HRP | Sigma | AP136P | Rat Polyclonal, IgG | 1:15000 |
| Anti-Goat IgG-HRP | Santa Cruz | sc-2354 | mouse monoclonal, IgG | 1:10000 |

**Supplementary Table 4.** Antibody list for immunohistochemistry.

| **Antibody** | **Vendor** | **Catalog #** | **Host Species** | **Dilution** |
| --- | --- | --- | --- | --- |
| Phosphorylated alpha synuclein | Cell Signaling Technology | 23706 | Rabbit Monoclonal, IgG | 1:500 |
| Anti-Iba1 | Fujifilm Wako | 019-19741 | Rabbit Polyclonal, IgG | 1:400 |
| CD68 | BioRad | MCA1957 | Rat Monoclonal, IgG2a | 1:200 |

**Supplementary Figure 1.**
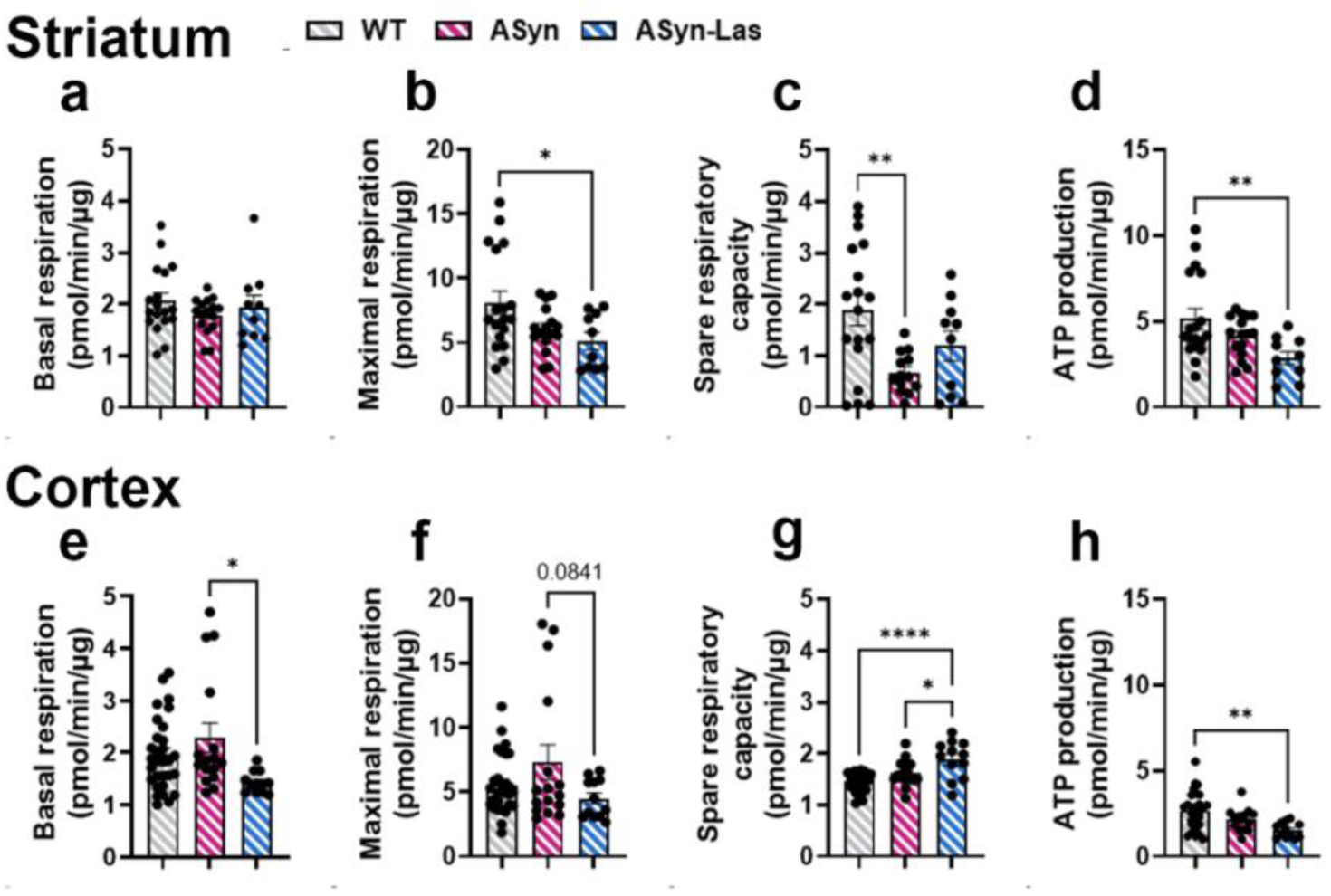
Oxidative phosphorylation changes in mitochondria of aged mice. WT mice are represented by gray bars with white diagonal stripes, ASyn mice by magenta bars with white diagonal stripes, ASyn-Las mice by blue bars with white diagonal stripes. Mitochondrial function in the striatum is represented in panels (a–d), and in the cortex in panels (e–h). Basal respiration is shown in (a) and (e), maximal respiration in (b) and (f), spare respiratory capacity in (c) and (g), and ATP production in (d) and (h). All values are presented as OCR (pmol/min/µg protein). Spare respiratory capacity is calculated as the difference between the maximal respiration after FCCP injection and basal respiration: Spare Respiratory Capacity = (Max rate after FCCP) – (Basal rate). *** p < 0.001, ** p < 0.01, * p < 0.05; One-way ANOVA with Tukey’s multiple comparisons test; mean ± SEM, n = 9–30 individual wells/group.

